# Learning from invariants predicts upcoming behavioral choice from spiking activity in monkey V1

**DOI:** 10.1101/2020.01.10.901504

**Authors:** Veronika Koren, Ariana R. Andrei, Ming Hu, Valentin Dragoi, Klaus Obermayer

## Abstract

Animals frequently make decisions based on sensory cues. In such a setting, the overlap in the information on the stimulus and on the choice is crucial for the formation of informed behavioral decisions. Yet, how the information on the stimulus and on the choice interact in the brain is poorly understood. Here, we study the representation of a binary decision variable in the primary visual cortex (V1) while macaque monkeys perform delayed match-to-sample task on naturalistic visual stimuli close to psychophysical threshold. Using population vectors, we demonstrate the overlap in decoding spaces on binary stimulus classes “match/non-match” and binary choices “same /different” of the animal. Leveraging this overlap, we use learning from the invariant information across the two classification problems to predict the choice of the animal as a time-dependent population signal. We show the importance of the across-neuron organization and the temporal structure of spike trains for the decision signal and suggest how noise correlations between neurons with similar decoding selectivity are helpful for the accumulation of the decision signal. Finally, we show that decision signal is primarily carried by bursting neurons in the superficial layers of the cortex.

**Author summary:** V1 is necessary for normal visual processing and is known to process features of visual stimuli such as orientation, but whether V1 also encodes behavioral decisions is an unresolved issue, with conflicting evidence. Here, we demonstrate that V1 encodes a mixed variable that contains the information about the stimulus as well as about the choice. We learn the structure of population responses in trials pertaining to the variable “stimulus+choice”, and apply the resulting population vectors to trials that differ only about the choice of the animal, but not about the stimulus class. Moreover, we learn structure of population responses on time-averaged data and then apply it on time-dependent (spiking) data. During the late phase of the trial, this procedure allows to predict the upcoming choice of the animal with a time-dependent population signal. The spiking signal of small neural population is sparse, and we hypothesize that positive correlations between neurons in the same decoding pool help the transmission of the decision-related information downstream. We find that noise correlations in the same decoding pool are significantly stronger than across coding pools, which corroborates our hypothesis on the benefit of noise correlations for the read-out of a time-dependent population signal.

## Introduction

Animals frequently make decisions based on the observation of the stimuli in their environment. Decision-making based on sensory information is traditionally studied with the measure of choice probability (CP) [1–10]. The CP measures the covariation between neural activity and the upcoming choice behavior of the animal. As we observe spike trains of single neurons, the CP is computed from distributions of spike counts that lead to a particular choice. If distributions are far apart, the choice can be predicted with high probability while highly overlapping distributions of spike counts give weak or non-significant CP. Weak but significantly better than chance CPs have been reported in a variety of sensory areas [1–5,11] including V2 [12]. The evidence on CP in V1 is disputed, since some studies found no evidence for CP in V1 [12], but the same authors subsequently found significant CPs in V1 during a coarse orientation-discrimination task with artificial stimuli [13], arguing that, in order to observe CP, neurons must be topologically organized with respect to the task feature. Weak CPs in V1 have been reported by other studies [9, 14], while robust choice signals have been reported in V1 when decoding from Local Field Potentials in a coarse orientation difference task with drifting gratings [10], as well as from spike counts of single neurons [8, 15], together demonstrating that the activity of neural populations in V1 does carry the information about the choice of the animal.

The hypothesis behind the CP framework is that it captures the choice related sensory evidence. As it is measured in single neurons, what informs the CP is the change in the spike count of the neuron, conditioned on the choice of the animal. Sensory neurons are tuned to features of sensory stimuli and within the CP framework, pooling the activity across neurons with a particular tuning corresponds to collecting the evidence about the sensory variable that leads to a decision signal and allows the brain to generate an appropriate behavioral response [1, 16]. As a simple example with a binary stimulus orientation “vertical” and “horizontal” and a corresponding binary choice of the animal, an increase in the spike count of neurons with the preference for the vertical orientation increases the probability of animal’s decision for “vertical” compared to “horizontal” while an increase in the spike count of neurons with preferred orientation for “horizontal” increases the probability for the choice “horizontal” compared to the choice “vertical”. As such, the CP framework is tightly linked to the notion of tuning of single neurons to the stimulus feature that is informative for the choice, and there has to be a positive correlation between the activity of tuned neurons and the choice. The CP can be though of as a “vote” of a particular neuron that is always consistent with its tuning. A recent study, however, has questioned the CPs by reporting a negative correlation between the activity of V1 neurons and animal’s choice [7], as neurons fired more vigorously when the animal chose the orientation that was the opposite to their preferred orientation. This result is in contradiction with the main hypothesis of the CP framework that predicts an increase in the spike count in neurons with preferred, and not with the anti-preferred orientation.

The CP framework is based on the assumption that single neurons carry behaviorally relevant signals while a body of theoretical insights and data analyses suggest that behaviorally relevant signals in the cortex are encoded by neural populations rather than by single neurons. Recent years have seen the revival and development of the idea that behaviorally relevant variables are encoded by collective dynamics of neural ensembles [17–19], and population codes seem to be insightful about the neural function in the auditory [20, 21], motor [22], prefrontal [23], and visual cortex [24]. In particular, distributed population codes have been suggested by the analysis of large neural populations in V1 [25]. Moreover, distributed population codes implemented as spiking neural networks have been demonstrated as optimal neural architectures by the theory [19]. If cortical networks operate with (distributed) population codes, a single neuron measure of choice-related signals, such as the CP, might only capture a limited amount of information about the choice.

Often, choice-related signals in sensory areas are studied in experimental paradigms where a sensory variable is changed in a continuous fashion, leading to a categorical decision variable with a limited number of options - typically two as the decision variable is binary. To better capture the relation between the representation of the stimulus and the choice in V1, we here impose a behavioral paradigm with a binary and categorical stimulus class instead of a continuous sensory variable. Previous work on decision-making with categorical variables suggested that such variables might be computed in a higher brain area, possibly through a comparison circuit [26], and represented in low-level areas as a contextual cue through a feedback signal areas [27–29]. Such feedback signals have been shown to be precisely timed [30] with the timing of feedback signals being crucial for perception [31, 32]. Since choice signals in V1 are likely caused by top-down projections from higher-order brain areas [8, 15], we reason that the neural representation of the choice is build in in real time to contribute to animal’s behavior as it unfolds. This evidence favors time-dependent models of decoding to measures that require averaging of spiking activity across time.

In the context of decision-making based on sensory cues, encoding of the sensory information and reading-out of the sensory information that is relevant for the choice behavior are two distinct, but connected computational problems. In particular, to generate choice signals that are informed by sensory evidence, there has to be an intersection of information between the stimulus and the choice [33]. While the activity of higher cortical areas might carry partially independent signals about the stimuli and the choice in order to dissociate these two variables [34], it is unlikely that V1 can afford the representation of the choice that is independent of the stimulus. Here, we hypothesize that, in face of a binary stimulus class, V1 can learn to implement perhaps the simplest case of intersection between the stimulus and the choice. We suggest that V1 neurons modify their synapses with high-level cortical areas and learn a set of decoding weights to represent the stimulus class and the correct choice of the animal. Following this hypothesis, we learn population decoding weights in trials with correct choice, where there is the information about the stimulus class and about animal’s correct choice. We then utilize these decoding weights to compute the population signal in trials that only differ about the choice, but not about the stimulus. Such a generalized learning scheme is based on the idea of learning from invariants [35], where intelligent agents are able to generalize the learned structure between distinct datasets with overlapping information, as to avoid the necessity to re-learn similar structures over and over. The generalized learning of decoding weights allows to predict the future choice of the animal with a time-dependent population read-out of spiking activity.

## Materials and methods

### Data and code availability

The analysis was done with Matlab R2019 (Mathworks). Code is available in a public GitHub repository: https://github.com/VeronikaKoren/transfer_learning

### Experimental model and subject details

All experiments were conducted in accordance with protocols approved by The Animal Welfare Committee (AWC) and the Institutional Animal Care and Use Committee (IACUC) for McGovern Medical School at The University of Texas Health Science Center at Houston (UTHealth), and met or exceeded the standards proposed by the National Institutes of Health’s Guide for the Care and Use of Laboratory Animals.

Two male rhesus macaques (*Macaca mulatta*; M1, 7 years old, 15kg; M2, 11 years old, 13kg) were used in this study. Animals were trained on a delayed match-to-sample task with visual stimuli. As the animal successfully fixated within the fixation area, this automatically triggered the trial. Two naturalistic images appeared consecutively, with a delay period in between. The target and the test stimulus were shown for 300 ms each, with a variable delay period in between (800 to 1000 ms). The identity of stimuli, complex naturalistic images depicting an outdoor scene, changed from one recording session to the other. Within the same session, the target and the test stimuli could be either the same (condition “match”) or else the test stimulus was rotated with respect to the target (condition “non-match”). The difference in orientation was such as to keep the performance stable at around 70 percent correct on non-matching stimuli, and ranged between 3 and 10 degrees. Animal subjects were trained to communicate their choice by holding the bar for the choice “different” and releasing the bar for “same”. For the trial to be valid, the behavioral response had to be communicated between 200 and 1200 ms after the offset of the test stimulus.

Multi-unit signal was measured with laminar arrays comprising 16 channels with 100 *μ*– meter spacing between adjacent contacts. Electrodes captured neuronal activity across the cortical depth, from superficial to deep layers. The multi-unit signal was spike sorted and we analyzed all cells that responded to the stimulus with at least a 4-fold increase of the firing rate with respect to the baseline. The data was collected in 20 recording sessions, which gave 160 neurons. We analyzed 3 conditions, “correct match” (CM), “correct non-match” (CNM) and “incorrect non-match” (INM), where “correct/incorrect” refers to the behavioral performance and “match/non-match” refers to the stimulus class. In condition “incorrect match”, there were not enough trials to perform the analysis. Table 1 reports the summary statistics on the number of trials (biological replicates) in each condition.

**Table 1.** The minimal, maximal, and average number of trials across recording sessions, for each condition.

| Condition | Nb. trials min. | Nb. trials max. | Nb trials average |
| --- | --- | --- | --- |
| CM | 51 | 290 | 118 |
| CNM | 42 | 230 | 107 |
| INM | 21 | 98 | 39 |

### Quantification and statistical analysis

#### Single neuron analysis

The spike train of a single neuron *n* in trial *j* is defined as a binary vector of zeros and ones,

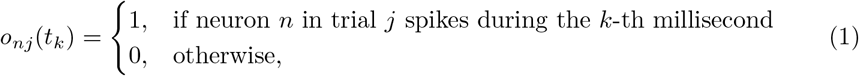

where *n* = 1,…, *N* is the neural index, *k* = 1,…, *K*, is the time index with step of 1 millisecond (ms), and *j* = 1,…, *J*_1_, *J*_1_ + 1,…, *J*_2_, *J*_2_ + 1,…*J* is the trial index. Trials were collected in conditions CM (*j* = 1,…, *J*_1_), CNM (*j* = *J*_1_ + 1,…, *J*_2_) and INM (*j* = *J*_2_ + 1,…, *J*).

The sensitivity index of the neuron *n* to the non-match of stimuli is defined as follows:

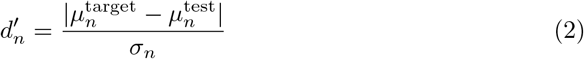

where 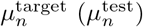 is the average spike count during the presentation of the target (test) stimulus in condition CNM, and *σ_n_* is the standard deviation of spike counts comprising trials from both target and test. To estimate the significance of *d*’, we randomly permuted (without repetition) trial labels for “target” and “test” and computed *d*’ with randomly labeled data *N*_perm_ = 1000 times. The p-value is then calculated by ranking the true *d*’ among the distribution of *N*_perm_ values of d’ computed with permuted labels. Here, as well as throughout the paper, random permutations are technical replicates [36].

#### Multivariate classification with Support Vector Machine

For multivariate classification, we use spike counts of neurons that were recorded in parallel. Spike counts were computed in the time window of [0, *K*] ms with respect to the onset of the stimulus (target or test), 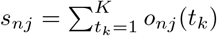. For each neuron, spike counts were z-scored across trials:

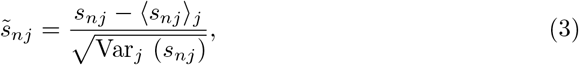

where 〈*s_nj_*〉_*j*_ and Var_*j*_ (*s_nj_*) are the mean and the variance of spike counts for the neuron *n* across trials from conditions that are used for classification.

Z-scored spike counts of *N* neurons recorded in parallel were utilized as input to the linear Support Vector Machine (SVM; [37]). Prediction accuracy of the classifier is computed as the balanced accuracy (*BAC*) on the hold-out test set:

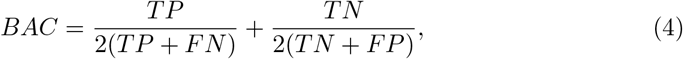

with *TP, TN, FP* and *FN* the number of true positive, true negative, false positive and false negative classifications, respectively. The performance measure of BAC corrects for the imbalance across conditions (see Table 1). The BAC is cross-validated *N*^CV^ = 100-times and reported values are averaged across cross-validations. The significance of the balanced accuracy is evaluated with the permutation test, utilizing *N*_perm_ = 1000 random permutations of class labels without repetition (see [38]).

When the population is divided in groups (see below), the prediction accuracy for the group of interest is computed by utilizing all simultaneously recorded neurons and removing the information from neurons that do not belong to the group of interest (see [39]). While class labels for the group of interest are correct, they are randomly assigned for the remaining of neurons, making those neurons contribute only chance-level signal. The advantage of this method is to keep the dimensionality of the dataset intact, thus avoiding the possible confound of imbalanced subpopulations. The significance of the difference in *BAC* between groups is tested with t-test on 100 instances of the *BAC* with random permutation of the class label (without repetition). The use of the t-test is justified as the distribution of *BAC* across random permutations is normal.

#### Estimation of the population vector

We estimate the population vector as the normalized vector of feature weights of the linear SVM. The population vector is computed using the activity of neurons recorded in parallel and refers to a specific binary classification problem.

One sample for the classifier is the vector of z-scored spike counts (eq. 3) of N simultaneously recorded neurons in trial *j*, 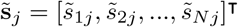. Linear SVM searches for an *N* – 1-dimensional plane (a hyperplane) that optimally separates data points [37] in conditions CNM and CM. The hyperplane is defined as

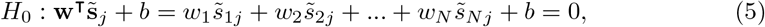

where **w** is the vector of feature weights and *b* is the offset of the hyperplane from the origin. On each side of *H*_0_, we can define a hyperplane that verifies the following:

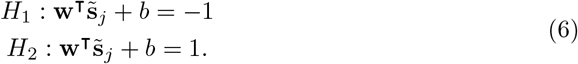

If the problem is linearly separable, all training samples verify the following the inequality:

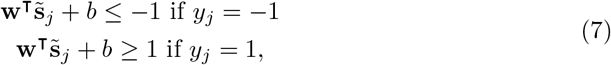

where *y_j_* ∈ { −1, 1} is the class label (*y_j_* = −1 in condition CNM and *y_j_* = 1 in condition CM). Training the linear SVM consists in maximizing the number of correctly classified samples and, at the same time, minimizing the distance between *H*_1_ and *H*_2_, which can be expressed with the Lagrangian as follows:

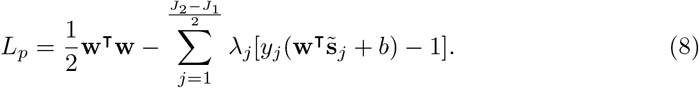

The first term on the right hand side of Eq (8) ensures the maximal distance between hyperplanes *H*_1_ and H_2_, and the second term on the right ensures correct classification. To find the minimum of the Lagrangian with respect to the vector **w**, we take the derivative of the Lagrangian with respect to **w** and set it to zero. This gives the expression for the vector of feature weights:

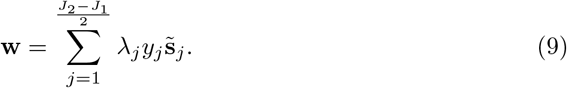

The vector of feature weights is normalized with the Euclidean (*L*^2^) norm,

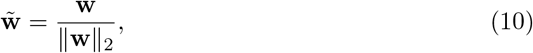

with 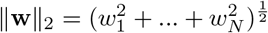. We refer to the vector 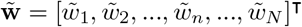 as the population vector and to the *n*–th entry of 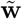, 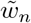, as the decoding weight of the neuron *n*.

#### Comparison of population vectors across classification problems

We estimate the population vector separately for classification problems on *stimulus* + *choice* and on *choice*, getting vectors 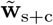 and 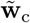. Classification of *stimulus* + *choice* utilizes trials from conditions CM and CNM, while classification of *choice* utilizes trials from conditions CNM and INM. The number of trials is imbalanced across conditions (namely, there are much less trials in condition INM compared to CM; see Table 1), and such imbalance can affect the population vector. We balance the number of trials with the bootstrap method. In each recording session, we find the number of trials of the condition with most trials, and randomly sample, with repetition, the same number of trials from the two other distributions. The bootstrap method applies only to comparison of population vectors, where reported results are averaged across bootstraps.

We measure the similarity of the population vectors 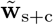 and 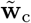 by computing the angle between them. From the dot product of vectors 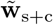 and 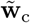,

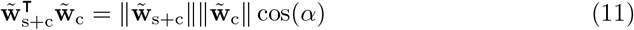

where *α* is the angle between the vectors, we compute the angle *α*:

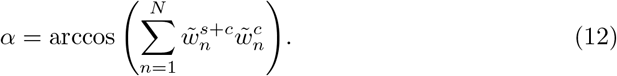

Notice that, since vectors are normalized, their Euclidean norm is unity, 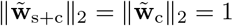. If the vectors 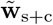 and 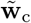 are similarly oriented, the angle between them is small. If, conversely, the orientation of two vectors is random between 0 and *π*, the angle between them is, on average, orthogonal (the average is across bootstrapped samples). The significance of the angle is evaluated with the permutation test. To construct the null model, we draw random vectors from the uniform distribution that have the same range and number of elements as the true population vectors, and compute the angle *α_p_* between the two random vectors. The p-value is computed by ranking the angle of true population vectors, *α*, among the distribution of *N*_perm_ angles of random vectors, *α_p_*. The test is significant if the following is true: 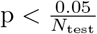, where the division with the number of tests implements the Bonferroni correction for multiple testing.

#### Population signal

Our decoding procedure comprises two steps, the estimation of the population vector (see the section “Estimation of the population vector”), and the computation of the population signal. These two steps implement an equivalent to the training and the validation step in a classical classification scheme. This is why the split of trials for the two steps is non-overlapping, such that no trials that have been used for the estimation of the population vector are used when computing the population signal.

The population vectors are estimated using all trials in condition CM and half of the trials in condition CNM (trials with index 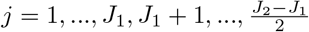). The population signal is computed on a hold-out set, utilizing the remaining half of the trials from condition CNM and all trials from condition INM (trials with index 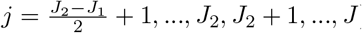). The split of trials in training and validation set in condition CNM is cross-validated with Monte-Carlo method, using *N_CV_* = 100 random splits. All reported results are averaged across cross-validations.

In the validation step, the population vector 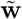 is utilized to compute the population signal as a weighted sum of spike trains. Consider the vector of spike trains of *N* simultaneously recorded neurons.

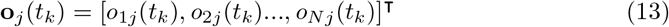

The population signal at time is the dot product of the vector of spike trains at time *t_k_* with the population vector (the latter does not change over time),

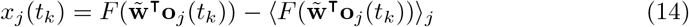

where 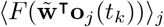 is the trial average, utilizing all trials of the validation set. By subtracting the trial average, we compute the deviation of the signal from the mean. Arguably, the deviation of the signal from its mean, rather than the absolute value of the signal, is a signal of biological relevance. As the transfer function, *F*(*y*(*t_k_*)), we use a convolution with an exponential filter,

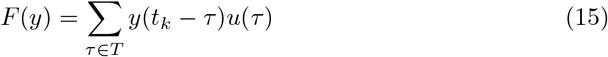

with *u*(*τ*) = *exp*(−*λτ*), *τ* ∈ *T*, with support *T* ms. The convolution with an exponential filter models the causal effect of the presynaptic spike on the neural membrane of the read-out neuron. The population signal *x_j_* (*t_k_*) is a time-resolved, low-dimensional representation of parallel spike trains in trial *j*.

Next, we average the choice signal across trials, distinguishing conditions CNM (decision “different”) and INM (decision “same”),

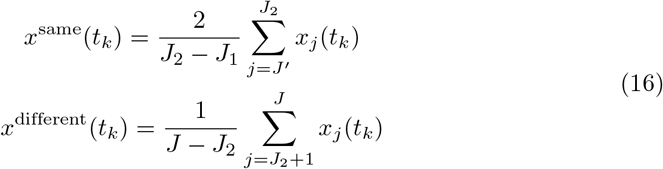

where 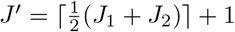, and [*z*] is the ceiling function. We obtain two time-dependent decision signals as in Eq 16 in every recording session. To have a more compact result, we compute the difference of the population signals in each recording session,

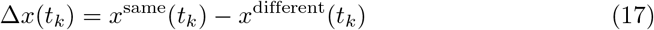

and average across sessions.

To assess the significance of the difference signal Δ*x*(*t_k_*), we use the permutation test, based on random permutation of class labels. We compute *N*_perm_ instances of the chance-level difference signal 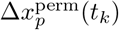, and compare the distribution of these values with the original difference signal Δ*x*(*t_k_*) in each timestep. When the difference signal of the true model, Δ*x*(*t_k_*), is outside of the distribution of results of the null model, 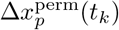 for *p* = 1,…, *N*_perm_, we consider that signals for *x*^same^(*t_k_*) and *x*^different^(*t_k_*) are significantly different.

#### Population signal with univariate decoding weights

To test decoding with univariate decoding weights, we replace population vectors from Eq 9 – 10 with weights that are computed from the area under the ROC curve of single neurons. The area under the ROC curve (AUC, [2]) is computed from distributions of spike counts of single neurons, distinguishing conditions CNM and CM. From the measure of AUC, we subtract the baseline, 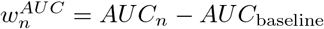, with *AUC*_baseline_ = 0.5. The weight 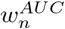 is positive if the neuron increased the spike count in condition CM compared to CNM, and negative if the neuron increased the spike count in condition CNM compared to CM. As we collect univariate decoding weights 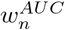 across neurons recorded on the same electrode, we get a vector 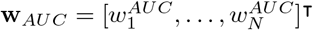. As with multivariate vector of decoding weights (see eq 10), we normalize the weight vector with the Euclidean norm,

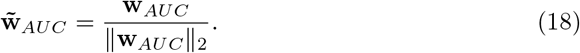

Normalized vector of univariate decoding weights 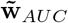 can now replace the population vector 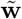 in eq 14 and be used to compute the population signal. With normalization in Eq 18, the length of the vector 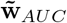 is unity, hence we can directly compare the population signals computed with multivariate and univariate decoding weights.

#### Criteria for division in subpopulations

We separate simultaneously recorded neurons into subpopulation with respect to the following four criteria: 1) sign and 2) strength of decoding weight, 3) burstiness and 4) location in the cortical layer. For criteria that take into account the population vector 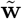 (i.e. the sign and strength of decoding weight), we utilize trials pertaining to the variable stimulus + choice (conditions CM and CNM). For all criteria, we used the length of the time window of *K* = 400 ms.

We separate the population into neurons with positive (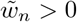; *plus* neurons) and negative decoding weight (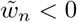; *minus* neurons) simply by considering the sign of the decoding weight of each neuron.

Neurons with strong and weak weight are distinguished by ranking the absolute value of decoding weight, 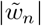, among the distribution of the same measure as decoding weights are computed with SVMs with permuted class labels. The strength of the weight of a single neuron *n* is a scalar, and the same result for models with permuted class labels is a distribution of *N*_perm_ values, where *N*_perm_ is the number of random permutations of class labels. If the strength of the weight of the neuron n is ranked within the first 25 % of weights from the null model, we assume that the neuron n has a strong weight, and we assume it has weak weight otherwise.

We distinguish bursting and non-bursting neurons utilizing a criterion based on the power spectrum (PS) of spike trains. It has been shown that bursting neurons have decreased PS in low and middle frequency ranges [40]. We compute the power spectrum of spike trains for every single neuron, using multiplication of the spike train with 5 Slepian tapers to increase the reliability of the estimation [41]. Power spectra are divided with neuron’s average firing rate, which, in case the spike train is a homogeneous Poisson process, gives a constant power spectrum of 1. We then compute the mean power spectrum for frequencies between 10 and 200 Hz, discarding frequencies below 10 Hz due to the short time window K. As a reference, the mean PS of a homogeneous Poisson process is 1, while the mean PS of a bursting neuron is typically below 1. We determine whether a neuron is bursting or non-bursting with a permutation test. We randomly permute N_perm_-times, without repetition, the spike timing of the neuron, and compute the distribution of mean PS from these data. We then rank the mean PS of the true model model among the distribution of corresponding results from spike trains with permuted spike timing. The neuron *n* is classified as bursting if its mean PS ranks within the lowest 5 percent of equivalent results of spike trains with permuted time index. If the neuron does not fulfill this criterion, it is classified as non-bursting.

Finally, we distinguish three cortical layers [42], the superficial (supragranular), the middle (granular) and the deep (infragranular) layer. The method for determining cortical layers utilizes the covariance matrix of the current source density and has been described previously (see [43]).

#### Choice-related signal in subpopulations

To gain better understanding of the information about the choice carried by subpopulations (as defined in the section “Criteria for division in subpopulations”), we compute population signals using only the information from a specific group of neurons that is of interest. The population signal of the group of interest is computed by removing the information about the choice in neurons that do not belong to the group of interest. This method ensures that the imbalance in the number of neuron across subpopulations does not influence the population signal. The information is removed by randomly permuting the class labels for “same” and “different” in the validation step, for neurons that do not belong to the group of interest. With permutation of the class label, the association between decoding weight of the neuron and the spike train is random, resulting in a signal that is close to zero at all times.

As an example, we detail this method for the case of decision signals in neurons with positive and negative decoding weights, but the same methods is used to compute the decision signal for subpopulations of neurons with strong and weak weights, for bursting and non-bursting subpopulations and for subpopulations in cortical layers.

The decision signal of *plus* neurons is computed with the spike train 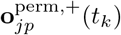, where the label of the spike train is correct for the *plus* neurons, and random (i.e., correct or incorrect with equal probability) for the *minus* neurons,

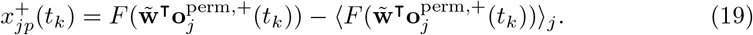

Next steps are as described previously (see Eq 16). Iterating this procedure *N*_perm_–times, with *p* = 1,…, *N*_perm_ random permutations of class labels for neurons that are not of interest, and then averaging across permutations, we get the population signal for *plus* neurons. Similarly, the population signal of *minus* neurons is computed by utilizing the spike train 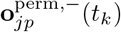, where the label of the spike train is now correct for the *minus* neurons, and random for the *plus* neurons.

#### Cross-correlation of population signals

The cross-correlation is used to measure the time-dependent relation of population signals between two subpopulations of simultaneously recorded neurons. Here we detail the procedure for computing the cross-correlation between population signals of *plus* and *minus* neurons, but the same procedure is used for bursting and non-bursting neurons and for neurons with weak and strong decoding weights.

The cross-correlation between population signals of *plus* and *minus* neurons in trial *j* and for the permutation instance *p* (see the section “Choice-related signal in subpopulations”) is:

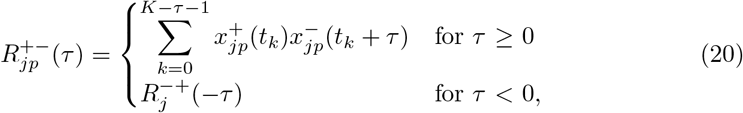

with time lag *τ* = 1, 2,…, 2*K* – 1. The cross-correlation is normalized with the autocorrelation at zero time lag,

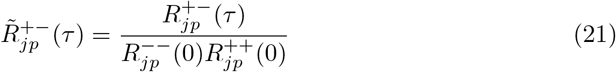

where *R*^++^ (*R*^−−^) is the autocorrelation function for *plus* (*minus*) neurons,

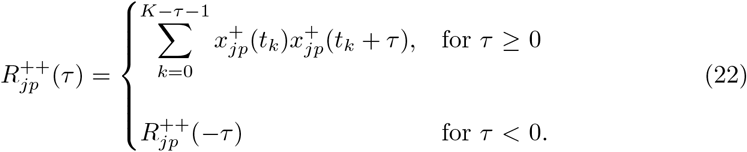

The correlation function in eq 21 is then averaged across trials and across permutations. Since there was no difference in the correlation across conditions, we used all trials from the validation set.

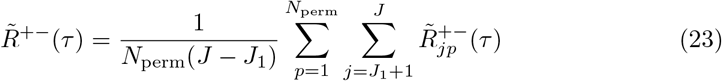

The significance of the cross-correlation in eq 23 is evaluated with the permutation test, where signals 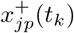 and 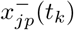 are computed with permutation of class labels. Additionally, we use random assignment to the group of *plus* and *minus* neurons in Eq 20.

The cross-correlation between population signals in cortical layers is computed by taking into account the population signals from two layers at a time, i.e.,

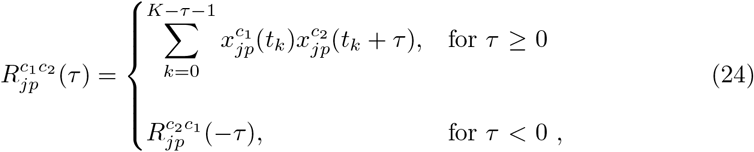

with (*c*_1_, *c*_2_) ∈ {(*SG*, *G*), (*SG*, *IG*), (*G*, *IG*)}. The rest of the procedure is the same as in Eq 21–23.

#### Noise correlation of spike timing

Let *n*, *m* ∈ {1,…, *N*} be fixed indices of two neurons, *n* ≠ *m*. We define the spike trains of neurons *n* and *m* in trial *j* by *f_j_*(*t_k_*):= *o_nj_*(*t_k_*) and *g_j_*(*t_k_*):= *o_mj_*(*t_k_*). The cross-correlation measures the co-occurence of spikes as a function of time lag *τ* between the two spike trains

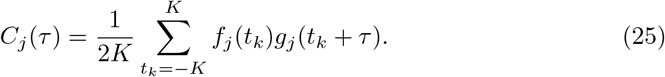

To make the cross-correlation independent of firing rates, we normalize it with autocorrelation of neurons *n* and *m* at zero time lag:

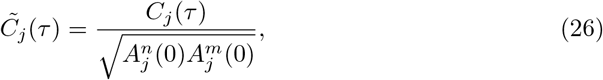

where the autocorrelation function is computed as follows:

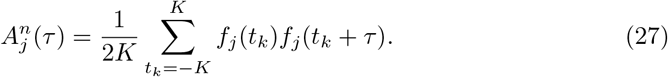

We now subtract the correlation that is attributed to trial-invariant processes:

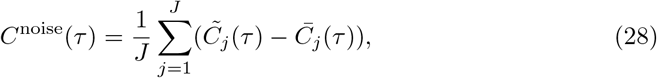

where the trial-invariant correlation function is computed by randomly permuting the order of trials of the spike train, without repetition, of one of the neurons:

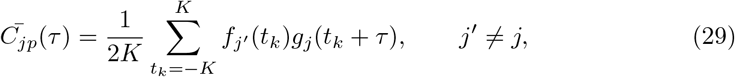

and 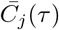 is the average of *N*_perm_ instances of random permutation. Finally, we sum the correlation function across the time lags [44]:

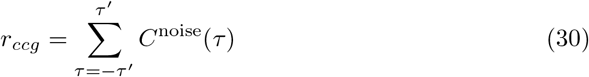

where *τ*’ is the maximal time lag.

To compute noise correlations, we use spike trains at the resolution of experimental recordings (Δ*t* =1 ms) and do not apply any further binning. When noise correlations are measured within a subpopulation of neurons, the entire procedure (Eq 25–29) applies to neuronal pairs from the specific subpopulation. The subpopulations are defined based on decoding weights from the classification problem *stimulus* + *choice*, while the correlation function, is computed using trials corresponding to the variable *choice*.

## Results

### Decoding spaces are similar between classification problems on *stimulus* + *choice* and *choice*

Two adult male macaques were trained on a delayed match-to-sample visual task. The subject visualized the target and the test stimuli, with a delay period in between (Fig 1A). The stimuli were complex naturalistic images in black and white, depicting an outdoor scene, and their identity changed from one recording session to another [45]. The target and test stimulus were either identical (condition match) or not (condition non-match), with the only possible difference being the change in the orientation of the test stimulus. In condition non-match, the angle of rotation of the test stimulus was random (clockwise or anti-clockwise), close to psychophysical threshold (between 3 and 12 deg) and was kept fixed throughout the recording session. The task of the subject was to communicate its decision about the stimulus class (“same” or “different”). The multiunit signal was captured with linear arrays, inserted perpendicularly to the cortical surface, recording the activity of neuronal populations across the cortical depth (see methods). Example spike rasters for three simultaneously recorded neurons, belonging to the superficial (supragranular or SG), middle (granular or G) and deep (infragranular or IG) cortical layer, are shown on Fig 1C. The difficulty of the task (i.e., the angle of rotation of the test stimulus) was calibrated during the experiment, as to keep the performance at 75% correct (Fig 1C).

**Fig 1.**
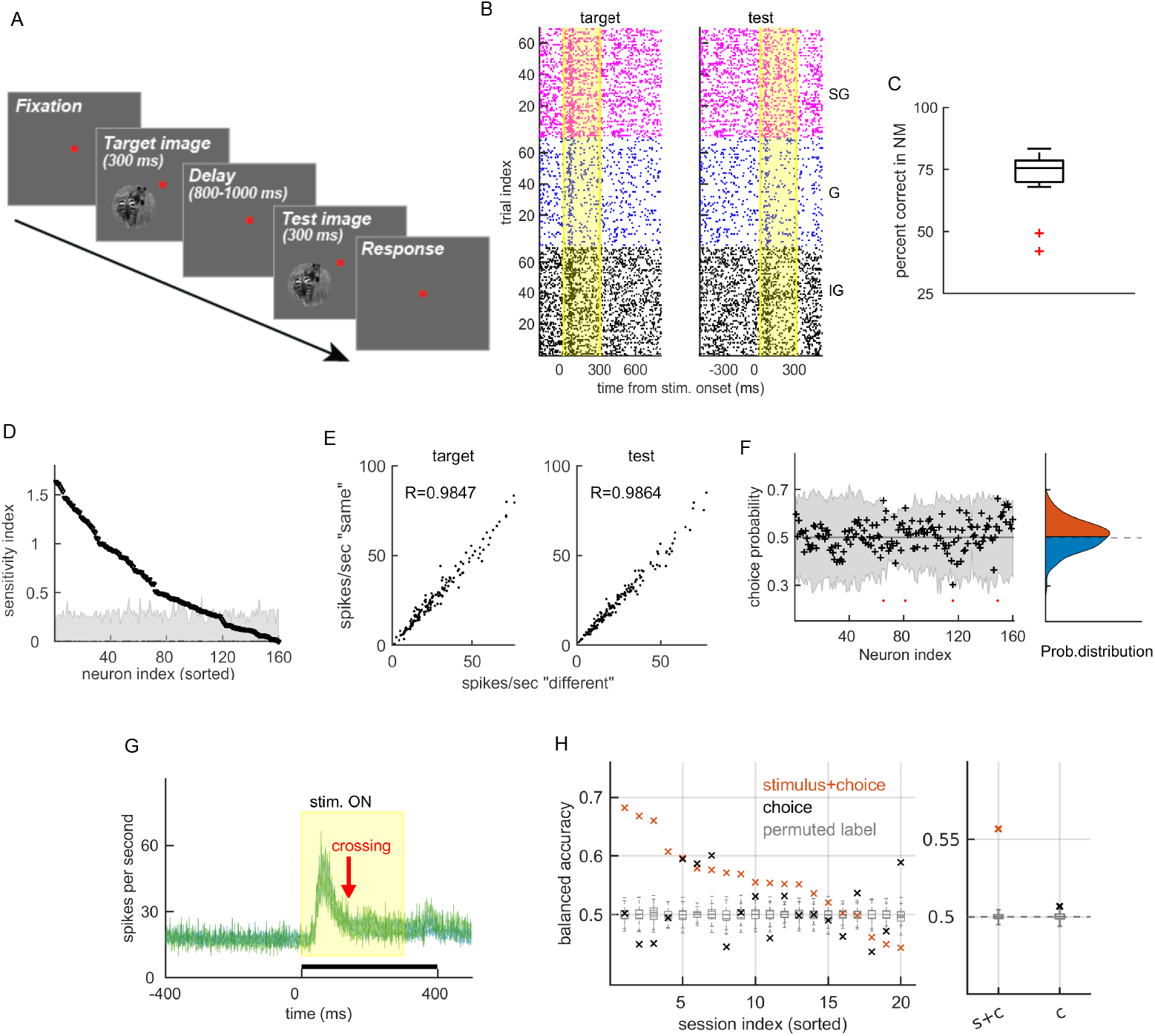
Single neuron statistics, PSTH and multivariate classifier do not predict animal’s choice. A: Experimental paradigm. During the experimental trial, a target and a test image consecutively appear on the screen, interleaved with a delay period. B: Spike rasters of three neurons from the same recording session centered on the onset of the target (left) and the test stimulus. Neurons are located in the superficial cortical layer (magenta), middle layer (blue) deep layer (black). The window when the stimulus is is on is marked by the yellow region. C: Percent correct trials on non-matching stimuli. D: Sensitivity index (*d*’) of single neurons to non-match of the stimuli, sorted from the strongest to the weakest. The gray area shows 95 % of results with permuted labels for “target” and “test”. E: Scatter plot of the mean firing rate for the choice “different” (x-axis) and “same” (y-axis). *R* is the Pearson correlation coefficient. F: Left: CP of single neurons (black), and the distribution of CPs for models with permuted class labels (gray). Right: Distribution of CPs from the left. G: Population Peri-Stimulus time histogram for choices “same” (green) and “different” (blue), with the mean ± SEM for the variability across sessions. The yellow window indicates the presence of the stimulus and the black bar indicates the time window we typically used for the analysis. The red arrow marks the time of the zero crossing of the decision signal in Fig. 3E. H: Prediction accuracy of the linear SVM for the classification problem on stimulus + choice (orange) and on choice (black). Left plot shows results in recording sessions and right plot shows session-averaged results. Gray boxplots show the distribution of results of models with permuted class labels. Parameter for D-H: Time window of *K* = [0, 400] ms with respect to the onset of the test stimulus (D-F,H). out of 160 neurons had a significant CP (permutation test). Counting also the neurons with non-significant CP, there was an excess of neurons with *CP* > 0.5 (*p* = 0.0140, permutation test), however, seen the weakness of this effect, the population CP did not seem to be a strong predictor of animal’s decision. The population Peri-Stimulus Time Histogram also did not allow to discriminate animal’s choice (Fig 1G).

Single neuron statistics and population averages are not predictive of the choice of the animal. Most single neurons were sensitive to the non-match of stimuli (Fig 1D), with the sensitivity index *d*’ (see methods) of 133 out of 160 neurons being significant to the change in orientation between the target and the test stimulus (permutation tests with permutation of class labels; all permutation tests in the paper use *N*_perm_ = 1000 permutations). Firing rates varied strongly across units, but not across the decision variable for the choices “same” vs. “different” (Fig 1E). CPs of single neurons were both bigger and smaller than the baseline (*CP*_baseline_ = 0.5), with an increase (*CP* > 0.5) as well as a decrease (*CP* < 0.5) in the firing rate with the choice “same” compared to the choice “different” (Fig 1F). After correction for multiple testing, only 4 Patterns of spike counts across the population predict the variable *stimulus* + *choice*, but not the variable *choice*. We applied a linear Support Vector Machine (SVM), the classifier with optimal generalization performance [46], on spike counts of populations of simultaneously recorded neurons during the presentation of the test stimulus. Prediction of the variable *choice* was not significantly different from chance (average balanced accuracy (BAC) of BAC = 0.507; not significant; permutation test with random permutations of class labels). Variable *stimulus* + *choice*, meanwhile, is predicted with better than chance accuracy (Fig 1H), demonstrating that patterns of spike counts are predictive of the choice when both choices are correct. The variable *stimulus* + *choice*, changes about the choice (“same”/”different”) as well as about the stimulus class (“match”/”non-match”; see Table 2), and from current results, one is tempted to conclude that the activity of neural populations in V1 contains the information about the stimulus class, but not about animal’s decision. However, the prediction of the stimulus class without the information about the choice is significantly lower than the prediction of the stimulus that is accompanied with correct choice (*p* < 0.001, permutation test; *BAC*^s+c^ = 0.557, *BAC*^S^ = 0.536), suggesting that the decision variable nevertheless has a significant impact on the neural activity in V1. The variable *choice* might not be possible to predict with classical decoding paradigms, because it comes with a mixture of correct and incorrect behavior (Table 2). When the choice is correct, we expect it to be mainly caused by correct inference on sensory information by the brain. Causes of an incorrect choice, however, can be many, for example, fluctuation of attention [47], failure of transmission of the sensory information downstream [45], noise in the feedforward input [48] or an error in the process of inference [49]. The coexistence of different causes in the same data can result in a chance-level prediction. Nevertheless, the choice of the animal can still be a latent variable that influences neural activity.

**Table 2.** Binary classification problems. The condition “incorrect match” has not been considered due to the insufficient number of trials.

| condition | stimulus | choice | variable |
| --- | --- | --- | --- |
| correct non-match | non-match | “different” | stimulus +<br>choice (s+c) |
| correct match | match | “same” |  |
| correct non-match | non-match | “different” | choice (C)<br>(+ performance) |
| incorrect non-match | non-match | “same” |  |
| correct match | match | “same” | stimulus (S)<br>(+ performance) |
| incorrect non-match | non-match | “same” |  |

If animal’s choice influences neural activity in the classification problem on *stimulus* + *choice*, we expect partially overlapping decoding spaces in the classification problems on *stimulus* + *choice* and on *choice* (see Table 2). Assuming a linear classification model, we use again the linear SVM to compare the orientation of linear separation boundaries (hyperplanes) across the two classification problems. Let us have *N* neurons observed simultaneously from which we collect data samples 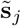, where 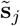 is a vector of z-scored spike counts of the *N* neurons in trial *j* (see methods). In general, the primal problem of the linear SVM can be expressed as follows:

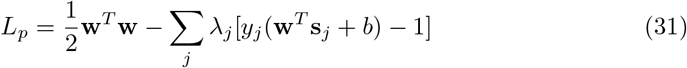

where **w** = [*w*_1_,…,*w_N_*]^T^ is the vector of feature weights, *λ_j_* and *y_j_* ∈ { −1, 1} are the Lagrange multiplier and the binary label in trial *j*, respectively, and *b* is the offset of the separation boundary from the origin. The optimal solution to the primal problem is a linear separation boundary in the *N*-dimensional space of neural activities that is optimal for separating binary classes “correct match” and “correct non-match”. We are interested in the orientation of the separation boundary, since the offset is not relevant for our problem. The offset would be relevant in case of a global change in the firing rate with the decision variable, which is not the case here (see Fig 1E-G). The orientation of the separation boundary in the *N*–dimensional space of neural activities is fully determined by the vector of feature weights **W**, with the separating boundary being perpendicular to the vector of feature weights. The expression for the vector **W** is given by minimizing the primal problem in Eq (31 with respect to the vector **w**. This gives:

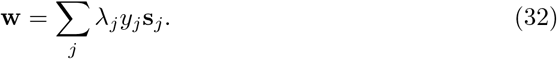

After normalizing the vector of feature weights with the Euclidean norm to keep the vectors in the same range across the recording sessions, we get the population vector 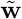:

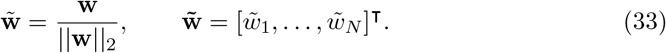

Note that normalization in Eq (33) does not change the orientation of the vector in Eq (32), but only normalizes its length. The *n*–th entry of the population vector 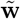, 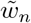, is the decoding weight of the neuron n, that summarizes the contribution of the neuron *n* to the separation boundary. A strong vs. weak decoding weight 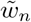 imply a strong vs. weak contribution of a particular neuron to the classification boundary. For example, if a neuron increases its spike count with respect to the baseline equally likely across conditions “correct match” and “correct non-match”, the activity of such a neuron is not helpful for the classifier and its decoding weight will be close to 0, but a neuron that consistently changes its spike count across conditions and is useful for classification will have a strong decoding weight. The sign of the weight, 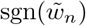, is informative about the increase/decrease of the spike count across conditions, and neurons with the same sign of the weight respond similarly to a particular condition.

We estimate the similarity of decoding spaces in classification problems on *stimulus* + *choice* and on *choice* by computing the angle between the population vectors. We separately estimate population vectors for classification problems *stimulus* + *choice* and *choice*, getting population vectors 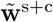 and 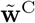 in each recording session. The angle between population vectors (Fig 2A) tells about the similarity of the orientation of separating boundaries in the two classification problems. If population vectors were independent, the angle between them would on average be orthogonal, *α*_independent_ = *π*/2, as it is when we compute population vectors from data with randomly permuted class labels (Fig 2B, gray boxplots). Deviation of the average angle from orthogonality means that the separation boundaries are partially overlapping. We find that the average angle between the population vectors deviates significantly from orthogonality, consistently for different time windows (*α* = [80.2, 80.8, 80.0] deg for the length of the time window of *K* = [300, 400, 500] ms, respectively; Fig 2B) and significant in all cases (*p* < 0.001, in all cases, permutation tests).

**Fig 2.**
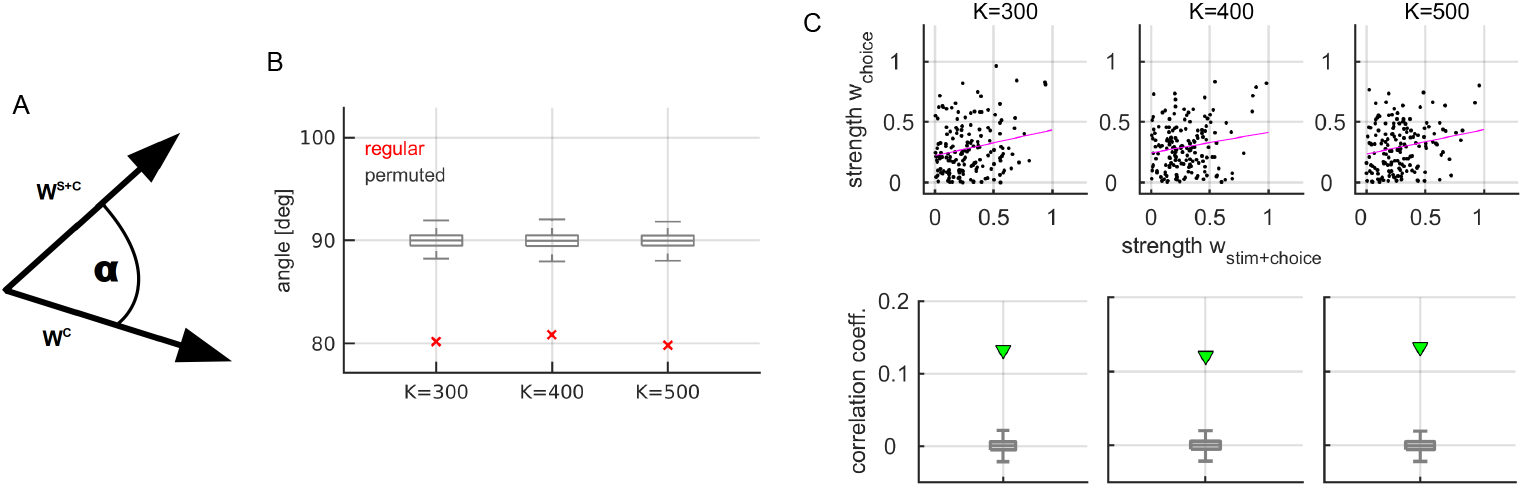
Similarity of decoding spaces in classification problems on *stimulus* + *choice* and on *choice*. A: Schema of the angle between population vectors in the context of *stimulus* + *choice* and *choice*. B: Average angle between population vectors (red cross) and the distribution of corresponding results by models with random weights (gray boxplots), for the length of the time window *K* = 300 (left), *K* = 400 (middle) and *K* = 500 ms (right). C: Top: Scatter plot of the absolute value of weights for the classification problem on *stimulus* + *choice* and on *choice*, utilizing the length of the time window of *K* = [300, 400, 500] ms. Bottom: Average correlation coefficient (green) for the data on the top. Gray boxplots show the distribution of corresponding results when decoding weights are computed with randomly permuted class labels. Parameter: *N*_perm_ = 1000

An alternative method to test the similarity of decoding spaces across the classification problems on *stimulus* + *choice* and on *choice* is to compare decoding weights of single neurons. To do so, we collect decoding weights across recording sessions, which gives two sets of decoding weights, 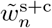 and 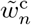, with neuron index *n* =1,…, *N*_total_ (*N*_total_ is the total number of neurons across recording sessions). We find a significant positive correlation (*r* = [0.13, 0.12, 0.13] across the two sets of weights, for the length of decoding window of *K* = [300, 400, 500] ms, respectively; Fig 2C). Positive correlation of decoding weights and similarity of population vectors across the two classification problems demonstrate similarity of decoding spaces in classification problems on *stimulus* + *choice* and on *choice*.

### Learning from invariants predicts the choice of the animal

After establishing the similarity of decoding spaces between classification problems on *stimulus* + *choice* and on *choice*, we exploit this similarity to learn from invariants and predict the variable *choice* as a time-resolved population signal. Learning from statistical invariants is a type of learning attributed to intelligent agents that goes beyond minimization of the expected error in a specific classification problem [35]. In intelligent systems, we expect learning to generalize on yet unseen but related problems. Learning from invariants allows an intelligent system to generalize a learned structure across distinct classification problems that share invariant statistical structure. In our case, classification problems on *stimulus* + *choice* and on *choice* both vary about the choice of the animal (Table 2), and have overlapping decoding spaces (Fig 2), hence, we hypothesize that learning on *stimulus* + *choice* could generalize over the variable *choice*. To implement this idea, we use decoding weights on the classification problem *stimulus* + *choice* (s+c), 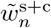, to weights parallel spike trains in trials that differ about the variable *choice* (c):

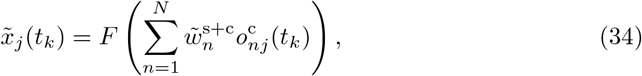

with *t_k_* the time index with resolution of the recorded spiking signal (Δ*t_k_* = 1 ms), *n* = 1,…, *N* the neural index for neurons recorded in parallel, *j* the trial index and *F*(*y*) = [*y* * *u*](*t*) the convolution with an exponential filter *u*(*τ*) = exp – *λτ*. From the absolute signal 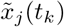 we subtract the running mean,

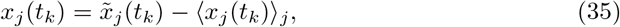

where 〈*x_j_*(*t_k_*)〉_*j*_ is the average across all trials. The time-dependent linear decoder in Eq (34–35) models the synaptic current at a hypothetical read-out neuron, generated by the spiking activity of the observed neural population. The convolution models the filtering of synaptic inputs at the membrane of the read-out neuron, with (*λ*)^-1^ the membrane time constant of the read-out neuron. Such a decoding procedure implements learning from invariants and differs from the regular training/validation scheme in two ways (Fig 3A). First, we learned the decoding weights in the classification problem that minimizes predicted error on the variable *stimulus* + *choice*, while we validated it on the variable *choice*. These are two related (Table 2) but nevertheless distinct classification problems. Second, we learned decoding weights on spike counts while we apply them on spike trains (Figure 3B). This is because decoding weights 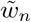 model changes in synaptic connections between observed neurons and the read-out neuron, a process that presumably takes place across days or weeks while the animal is training to perform the behavioral task. The signal about the choice, on the other hand, is presumably read-out by the brain in real time.

**Fig 3.**
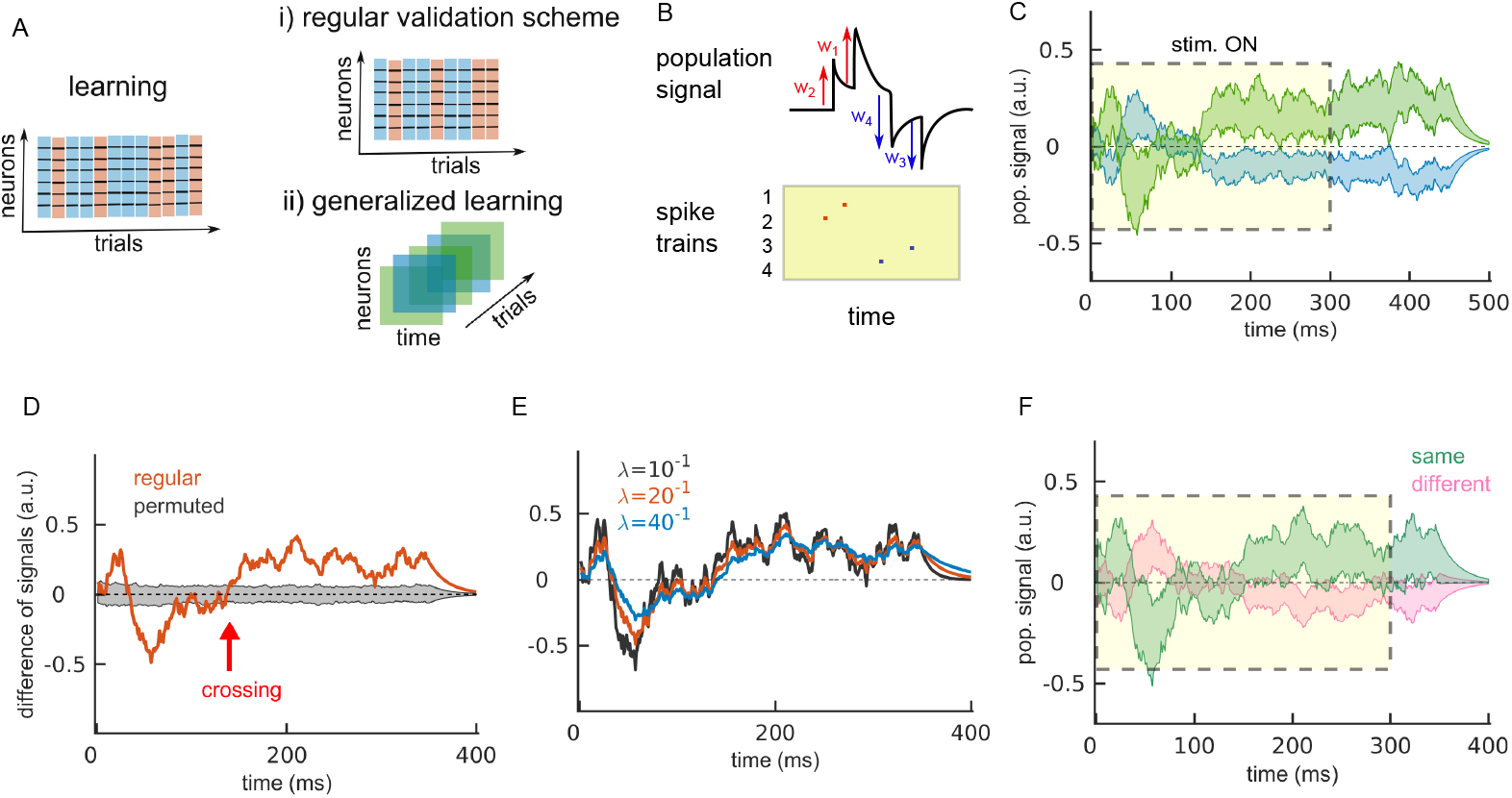
Learning from invariants allows the prediction of the upcoming choice. A: Learning from invariants (schematic). In the learning phase, the classifier learns from samples of parallel spike trains (left). Following the regular validation scheme, the same type of data in the same conditions would be used for validating the predictive power of the model on a hold-out set (top right). Here, we test learning from invariants (bottom right), where the training and the test set differ in the time scale of the data (from spike counts in the training set to spike trains in the validation set) and the classification problems (stimulus + choice in the training set and choice in the validation set). B: Computation of time-dependent population signal by weighting spikes with decoding weights (schematic). Every spike causes a jump of the population signal, followed by an exponential decay towards 0. Spikes of neurons with positive weights push the signal up and spikes of neurons with negative weights pull the signal down. C: The population signal for the decision “same” (green) and “different” (blue) with learning from invariants. We show the mean ± SEM for the variability across recording sessions. Results are cross-validated with 100 cross-validations and the learning and validation set are non-overlapping (see methods). D: Session-averaged difference of population signals for the regular model (red) and the model with permuted class labels (gray). E: Difference of decision signals for three values of the parameter *λ*. The exponential kernel used for convolution is normalized such that the area under the kernel is 1. F: Same as in C, but replacing decoding weights from Eq 32 with centered and normalized area under the ROC curve. Parameters: *λ*^-1^ = 20 ms, time window for C is [0, 500] ms, time window for D-F is [0, 400] ms.

With learning from invariants, a time-dependent decision variable can be predicted during the majority of the trial. Averaging the population signals in Eq 35 across trials that lead the same decision, we get distinct signals for animal’s choice “same” vs. “different” that persists even after 150 ms after the *offset* of the stimulus (Fig 3C). Since the variable *choice* does not *per se* vary about the stimulus class (both conditions in the variable *choice* are with non-matching stimuli; see Table 2), the population signal computed in Eq 34–35 reflects the information about the choice of the animal. To estimate the significance of the difference of the population signals pertaining to decisions “same” and “different”, we compute the difference of trial-averaged signals:

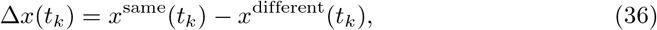

and compare it to same results computed with permutations of class labels (see methods). The difference signal Δ*x*(*t_k_*) is significant during the majority of the trial (Fig 3D), and the read-out of animal’s decision is stable starting from 140 ms after the onset of the stimulus (Fig 3C-D). Feedforward currents are expected to drive neural activity at the stimulus onset, while top-down and lateral inputs are expected to dominate the neural activity during the late phase of the trial [50]. The timing of 140 ms after the stimulus onset coincides with the end of transient firing at the stimulus onset (Fig 1G, red arrow), suggesting the relevance of top-down inputs, as opposed to bottom-up inputs, for the decision signal in the present setting. Note that the variable *choice* contains correct and incorrect behavior (Table 2), which is a likely explanation for unstable representation of the choice at the beginning of the trial. Properties of the time-dependent population signal decoded from the spiking activity are compatible with an internally generated decision variable resulting from a top-down projection from another cortical area.

Prediction of animal’s choice is robust to the time window used for decoding (S1 Fig), the time constant of convolution (Fig 3E), as well as the method for estimating decoding weights. As we replace decoding weights in Eq 31–33 with centered and normalized Area Under the ROC curve that we estimated for each neuron (see methods) the choice of the animal can still be predicted, even though less accurately (Fig 3F). This shows that optimal decoding weights estimated with the linear SVM are not required to predict the choice. Learning on trials with correct behavior (i.e., *stimulus* + *choice*), however, is necessary for prediction of the *choice*. When the learning and the validation step both utilize the variable *choice* (instead of learning on *stimulus* + *choice*), the resulting signals are not significant (S2 Fig). As expected, prediction of the choice during the visualization of target stimulus is at chance (S3 Fig), because during the presentation of the target stimulus, the information necessary for discrimination of matching from non-matching stimuli is not yet available to the visual system.

The correct sign of decoding weights, 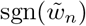, is necessary for predicting animal’s decision. The sign as well as the amplitude of decoding weights determine the orientation of the population vector in the N–dimensional space of neural activities. However, flipping the sign of the weight typically causes a bigger change in orientation than perturbing the amplitude (or strength) of the weight (Fig 4A, left). To test the importance of the sign and the amplitude of decoding weights, we replace the correct sign and amplitude with values randomly drawn from the uniform distribution. As we randomize the sign of decoding weights, prediction of animal’s decision is no longer possible, while using random amplitude still allows the predict animal’s decision (Fig 4B), even though the decision signal is now weaker with respect to the unperturbed signal. Further on, we compute the decision signals separately in subpopulations with positive and negative sign of decoding weight and estimate their time-lagged cross-correlation in the same trial (see methods). The decision signals of subpopulations with positive and negative weights are negatively correlated (Fig 4C), with half-width at the trough value at 18 ms (using the membrane time constant of the read-out unit of *λ*^-1^ = 20 ms). Negative correlation suggests competition between these subpopulations.

**Fig 4.**
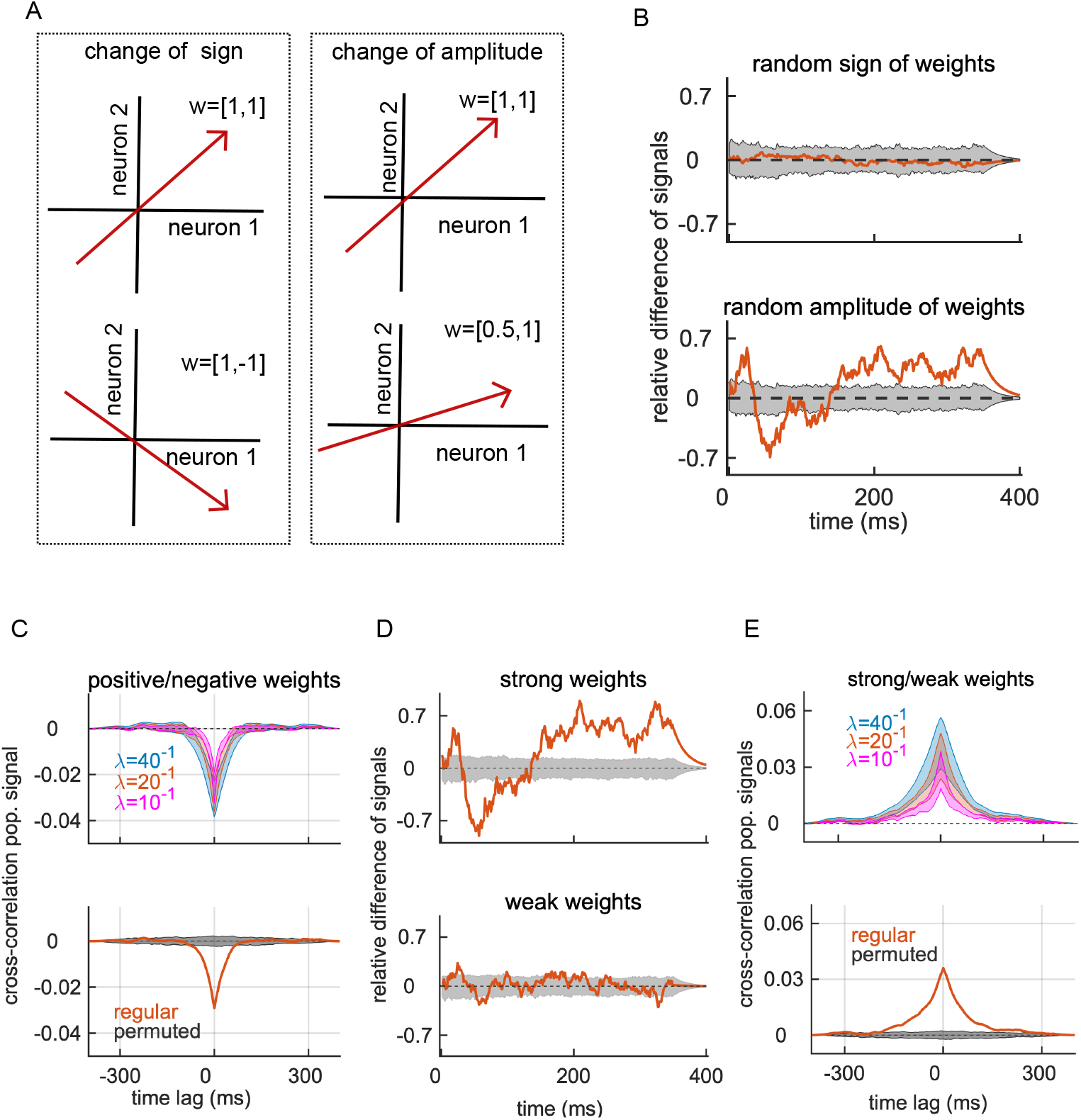
Correct sign of decoding weights is necessary for prediction of the choice. A: Effect of changing the sign of a decoding weight (left) and the amplitude of a decoding weight (right) on the orientation of the population vector in the space of inputs of N=2 neurons (schematic). Both types of perturbation change the orientation of the population vector (red arrow), but flipping the sign typically induces bigger changes. B: The difference of decision signals computed with random sign of weights (top) and random amplitude of weights (bottom). The gray area marks the distribution of models with permuted class labels. Signals are scaled with respect to the unperturbed signal (in Fig 3C) at maximum deviation from 0 (211 ms after the stimulus onset). C: Top: Cross-correlation of decision signals between neurons with positive and negative weights, for different values of the inverse time constant of convolution *λ*). We show the mean ± SEM across recording sessions. Bottom: Session-averaged cross-correlation for the model with *λ*^-1^ =20 ms (red) and the distribution of results for models with permuted class labels (gray area). D: Same as in B, but using the information from neurons with strong weights (top) and with weak weights (bottom). E: Same as in C, but for subpopulations of neurons with strong and weak weights. Parameters (when applicable): *λ*^-1^ = 20 ms, *N*_perm_ = 1000, Δ*t* = 1 ms.

The decision signal relies on the activity of a subpopulation of informative neurons. We split the population of simultaneously observed neurons in two groups with respect to the amplitude (or strength) of the decoding weight 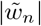, distinguishing neurons with strong and weak decoding weights (see methods). The groups are approximately balanced, with 79 and 81 neurons with strong and weak weights, respectively (as we collect the weights across recording sessions). As it follows from the definition of the linear SVM (see Eq 31–32), neurons with strong decoding weights are those that contribute the most to the population vector, while neurons with weak decoding weights contribute little or nothing. The subpopulation with strong decoding weights carries the decision signal, while the decision signal of the subpopulation with weak weights does not reach significance (Fig 4D). The cross-correlation between decision signals of subpopulations with strong and weak weights is positive (Fig 4E), with half-width of the peak value at 39 ms (using the membrane time constant of *λ*^-1^ = 20 ms).

### Positive noise correlations between neurons from the same coding pool strengthen the time-dependent decision signal

An unresolved question in neuroscience is the role of noise correlations for neural coding. Positive noise correlations among neurons with similar tuning harm the quantity of information that a neural population transmits with a time-averaged signal [51]. In a population code based on time-averaged neural activity, one would therefore expect that noise correlations among neuronal pairs with similar tuning are weak, or at least weaker than noise correlations among pairs with different or opposite tuning. In contrast to this prediction of a theoretical model, positive interactions between signal and noise correlations in V1 have been observed in experiments [52]. Moreover, if positive correlations among neurons with similar selectivity were detrimental to the neural code, we would expect them to be weaker with correct compared to incorrect behavior. In contrast, stronger noise correlations [53] and across-neuron coordination [45] were reported in correct compared to incorrect behavior. This evidence suggest that, in some cases at least, noise correlations among neurons with similar selectivity might be useful for the read-out. While across-neuron noise correlations might harm the transmission of information when reading-out from samples of time-averaged neural activity (i.e., parallel spike counts), this is not necessarily the case when reading out from a time-dependent signal (i.e., parallel spike trains). A spike train of a single neuron is a signal that is transient and sparse in time. The presynaptic spike has an impact on the membrane potential of the read-out neuron for the duration of one membrane time constant of the read-out neuron, which is around 10-20 ms. Due to the sparsity and the transient effect of spikes on the postsynaptic neurons, the beneficial relation between neuronal selectivity and noise correlations in a time-dependent read-out is reversed with respect to the time-averaged read-out.

With a time-dependent read-out as in Eq (34), positive noise correlations among neurons with similar selectivity and negative noise correlations between neurons with different selectivity strengthen the population signal and increase its impact at the read-out neuron(s). In our case with a binary classification problem, we therefore expect neuronal pairs with the same sign of decoding weight (same decoding pool) to have stronger noise correlations than pairs with the opposite sign of decoding weight (opposite decoding pool). This organization of noise correlations is indeed observed in the data (Fig 5A,B). Then, due to normalization in Eq (35) where we subtracted the baseline of the neural signals, we get positive effective interaction between neurons from the same pool, and negative effective interaction across neurons from the opposite pools. Such spatio-temporal organization of spiking allows the population signal to accumulate over time and raise above the noisy background (Fig 5C, top), which improves detection of the population signal at the read-out neuron. If, on the contrary, neurons from the opposite pools fire in a random sequence, the population signal remains close to zero and can only convey noise to the read-out neuron (Fig 5C, bottom).

**Fig 5.**
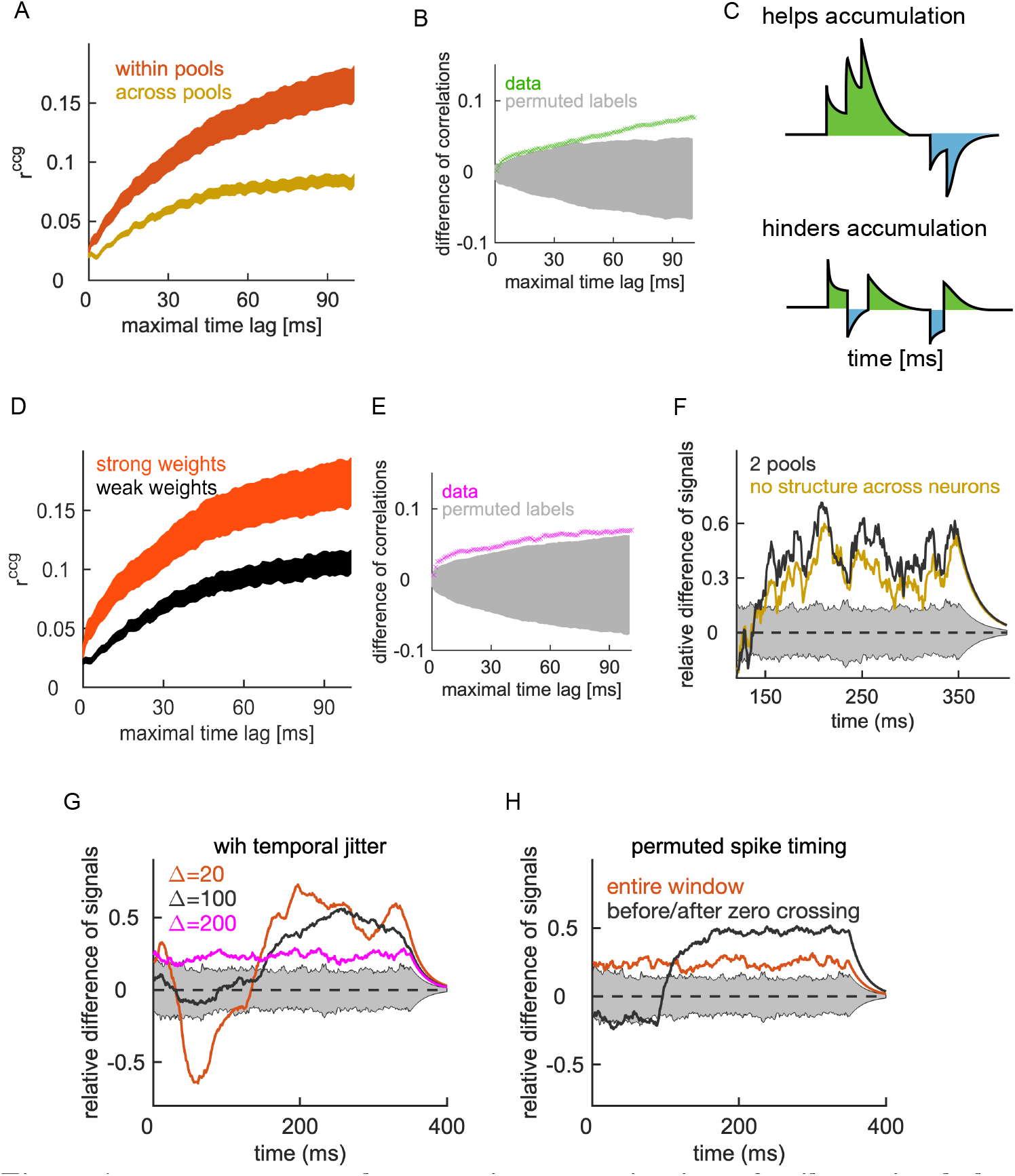
Across-neurons and across-time organization of spike trains helps the transmission of the decision signal. A: Noise correlation of spike timing for neuronal pairs with the same sign of the weight (within pools; orange) and with the opposite sign of the weight (across pools; ochre). Noise correlations are computed as the area under the cross-correlogram (*r^ccg^*), and the x-axis is the maximal time lag used for computing the area. We show the mean *r^ccg^*± SEM for the variability across pairs (*N*^within^ = 207,*N*^across^ = 514). B: Difference of average noise correlation (*r*^within^ – *r*^across^; green) and the distribution of corresponding results results with *N*_perm_ random permutations of labels for label {“within”, “across”}. C: Schematic of spike patterns that either help or hinder the accumulation of a time-resolved population signal. D: Same as in A, but showing noise correlations among neuronal pairs with strong weights (orange) and weak weights (black). E: Same as in B, but showing the difference of average noise correlation (*r*^strong^ – *r*^weak^; black) and the distribution of results for *N*_perm_ random permutations of labels for labels { “strong”, “weak” } in gray. F: Difference of decision signals with permuted spikes across neurons. We permute spikes such as to maintain the coarse organization in 2 pools of neurons with positive and negative sign of the weight (black), and across all neurons (ochre). The gray area is for the distribution of signals with permuted class labels (the null model). The difference signals is scaled with respect to the unperturbed signal at maximum deviation from 0 (i.e., the decision signal in Fig 3C at 211 ms after the stimulus onset). We permuted neuronal indices, randomly in each cross-validation cycle. G: Same as in F, showing decision signals with temporal jitter in the spike timing. H: Same as in F, showing the decision signal with permuted spike timing over the entire time window ([0,400] ms) or in two time windows before and after the zero crossing at 140 ms after the stimulus onset ([0,140],[141,400] ms). Parameters: time window for measuring correlations (A,B,D,E) and the effect of across-neurons structure is [140, 500] ms, time window for G and H is [0, 400] ms after the onset of the test stimulus; *λ*^-1^ = 20 ms, *N*_perm_ = 1000.

If noise correlations are helpful for the information transfer, we expect them to be stronger within subpopulations of neurons that are particularly informative for the task. By definition of the decoding model, neurons that are particularly informative for the task are neurons with strong decoding weights and neurons that are less informative have weak decoding weights (see Fig 4D). We therefore compare noise correlations among neuronal pairs with strong weights and weak weights. In line with our hypothesis on the benefit of noise correlations for the read-out of the time-dependent population signal, neurons with strong decoding weights show stronger noise correlations of spike timing than neurons with weak weights (Fig 5D,E).

Furthermore, across-neuron structure of spike trains is beneficial for the read-out of the decision signal. The benefit of noise correlations for coding suggested by results above (Fig 5A,D) implies that across-neuron and across-time organization in parallel spike trains carries information about animal’s decision. We test the effect of across-neuron structure in spike trains on the decision signal by removing this structure from spike trains, computing the decision signal on such perturbed data, and comparing the perturbed decision signal with the unperturbed one. First, we keep the coarse organization in two neuronal pools but selectively remove across-neuron structure within neuronal pools. Removing the structure within pools weakens the decision signal for about 31 % with respect to the original signal (Fig 5F, black). Next, we perturb the structure across all neurons. This further weakens the decision signal for another 9 %. In total, the across-neuron organization of spike trains therefore contributes a substantial 40 % to the read-out of the decision signal. This contribution gets partitioned into about 31 % for the organization in two neuronal pools (following the sign of decoding weight) and in 9 % for the fine organization within pools.

Further on, we test the effect of fine temporal structure of spike trains on the decision signal. Do do so, we jitter the timing of every spike within the time window of (± [20,100,200] ms) and then recompute the decision signal. The longer the jittering window, the weaker the decision signal (Fig 5G). More precisely, a jitter of 20,100 and 200 ms reduce the decision signal with respect to the signal with intact spike timing for 34, 57 and 75 %, respectively, measuring the difference at the maximal deviation of the original signal from zero. Note that the decision signal with 200 ms jittering window is close to losing significance. Even short jittering window of 5 and 10 ms reduce the decision signal for 16 and 22 %, respectively. We now test an even stronger perturbation of the temporal structure of spike trains on the read-out and permute the spike timing (for every neuron independently) over the entire time window between [0,400] ms after the stimulus onset. This weakens the decision signal for 79 % with respect to the decision signal with intact spike timing, and brings it close to noise (Fig 5F, red). We reason that a strong effect of shuffling the spike timing over a long time window might be partially due to different sources of the decision signal in the initial and late phase of the trial (see Fig 3D). If this is so, limiting the permutation to the early and late time window could improve the read-out. Indeed, limiting the permutation of the spike timing to before and after 140 ms after the onset of the test stimulus, we get a stronger decision signal (Fig 5F, black), that is however still 55 % weaker than the original signal.

### Activity of bursting neurons and of neurons in the superficial cortical layer is particularly informative about the choice of the animal

In the present setting, the read-out of the decision signal relies on decoding weights 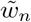 that have here been learned with a supervised learning method (using the linear SVM). In a biological network, learning of decoding weights requires plasticity of synapses between V1 neurons and a downstream brain area that is providing the top-down input. A recent study demonstrated a biologically relevant mechanism of synaptic plasticity that operates across the brain hierarchy [54], where learning of synaptic weights between a low- and high-level brain area critically relies on bursting of neurons in the low-level brain area. Seen these results, we hypothesize that bursting neurons might be important for the read-out of the choice signal in the present setting. We divide neurons into bursting and non-bursting, based on characteristics of the power spectrum (PS) of spike trains. The PS of bursting neurons are characterized by a reduced power in middle range frequencies [40] (Fig 6A), and we capture this effect by computing the area under the PS of the spike train and compare it with the same result as the spike train has been randomly permuted over time. A random permutation of the spike train results in a Poisson process, since each timestep is now equally likely to contain a spike. We classify a neuron as bursting if it has the area under PS smaller than the lower bound of models with permuted spike timing (Fig 6B; see methods), and as non-bursting otherwise.

**Fig 6.**
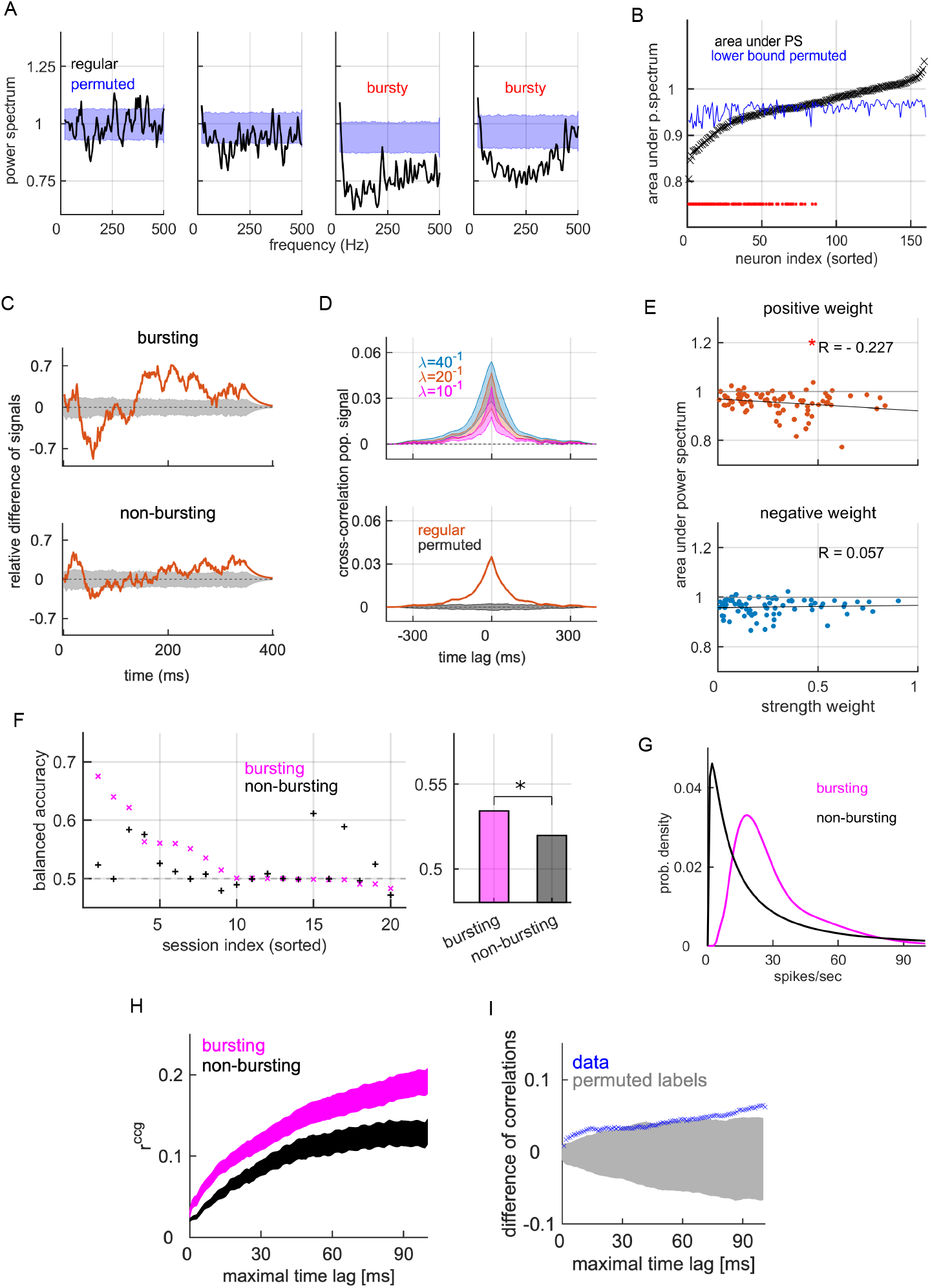
Bursting neurons carry more information about the choice than non-bursting neurons. A: Power spectrum for 4 representative neurons from the same recording session. We show the power spectrum for the true model (black) and the distribution of power spectra for models with randomly permuted spike timing (blue). Bursting neurons are those with reduced power in lower and medium range frequencies (middle right and right). B: Area under the power spectrum for all neurons, sorted from the lowest to the highest value (black), and the lower bound of the area under the power spectrum for models with permuted spike timing (blue). Red asterisk marks significance (permutation test, *α* = 0.05). There are 68 bursting and 92 non-bursting neurons. C: The difference of mean decision signals for bursting (top) and non-bursting subpopulations (bottom). Corresponding results of models with permuted class labels are shown as the gray area. D: Top: Cross-correlation of decision signals of bursting and non-bursting subpopulations. We show the mean ± SEM for the variability across recording sessions. Bottom: Session-averaged cross-correlation for the regular model (red) and the distribution of results of models with permuted class labels (gray area). E: Scatter plot of the area under the power spectrum vs. strength of decoding weight, distinguishing neurons with positive weights (top) and neurons with negative weights (bottom). R is the Pearson correlation coefficient. F: Balanced accuracy of the linear SVM on the classification problem stimulus + choice, using the information from bursting (magenta) or non-bursting subpopulations (black). G: Distribution of average firing rates for bursting and non-bursting neurons. H: Noise correlations of spike timing among bursting (N=198 pairs) and non-bursting neurons (N=231 pairs). We show the mean ± SEM for variability across pairs. I: Average difference of noise correlations (*r*^bursting^ – *r*^non-bursting^; blue) as a function of the maximal time lag, and the distribution of corresponding results of models with random assignment of the label {“bursting”, “non-bursting”}). Parameters: *λ*^-1^ = 20 ms (C), time window is [0, 400] ms after the onset of the test stimulus for B-G, and [140,500] ms for H-I; *N*_perm_ = 1000.

As we now compute the decision signal for bursting and non-bursting subpopulations, we find that the decision signal in bursting subpopulations is clearly stronger than in non-bursting subpopulations (Fig 6C). Decision signals of bursting and non-bursting subpopulations are positively correlated and have a peak at zero time-lag (Fig 6D). Importantly, for neurons with positive weight 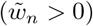, the strength of decoding weight is negatively correlated with the area under the PS (Fig 6E), meaning that the more bursting the neuron, the stronger decoding weight it tends to have. The correlation between the strength of the decoding weight and the are area under PS is significant for neurons with positive weights, but not for neurons with negative weights (*p* = 0.012 for neurons with positive weights, *p* = 0.610 for neurons with negative weight; permutation test with 1000 random assignment of labels {“bursting”, “non-bursting”}).

Since bursting is related to learning of synaptic weights, we reason that bursting neurons might be better than non-bursting neurons also in predicting stimulus + choice with time-averaged data. Using again the linear SVM on parallel spike counts, but this time separating the information carried by bursting and non-bursting subpopulations, we find that subpopulations of bursting neurons predict the variable stimulus + choice better than the non-bursting neurons (*p* < 10^-17^, two-tailed Wilcoxon signed-rank test; Fig 6F). This shows that the *time-averaged* activity in bursting neurons is more informative about stimulus + choice than in non-bursting neurons. Even though the absolute firing rate does not inform the classifier in Fig 6F, it is noteworthy that bursting neurons have substantially stronger firing rates than non-bursting neurons (Fig 6G). Furthermore, consistent with our hypothesis on the benefit of noise correlations for a time-dependent population read-out, bursting neurons have markedly stronger noise correlations of spike timing compared to non-bursting neurons (Fig 6H-I).

At last, we divide neurons with respect to their location along the cortical depth in three cortical layers. Cortical layers differ in the source of their inputs, as the middle layer receives the strongest feed-forward input, while the superficial layer has been suggested as a likely target of top-down inputs [50, 55]. Moreover, cortical layers were shown to differ in their correlation structure [42], which has direct consequences on the population code used here (see Fig 5C). We distinguish the superficial layer (N=48), the middle layer (N=51) and the deep layer of the cortex (N=61), using a method based on the current source density [43]. As we compute the decision signal in each layer, we find substantial and qualitative differences across layers (Fig 7A), and in particular, the strongest decision signal is carried by the superficial layer. Decision signals are weakly positively correlated across all pairs of layers, and cross-correlations peak at 0 time-lag in all cases (Fig 7B). If the decision signal originated from a bottom-up input, we would expect to see a significant decision signal in the middle layer, with potential decision signals in other layers following after a time delay. The fact that the decision signal in the middle layer is weak, but strong in the superficial layer is in line with our hypothesis about the top-down origin of the decision signal. Moreover, the fact that the cross-correlation of decision signals has a peak at zero time lag indicates that decision signals originate from a common (top-down) input rather than from interaction across cortical layers.

**Fig 7.**
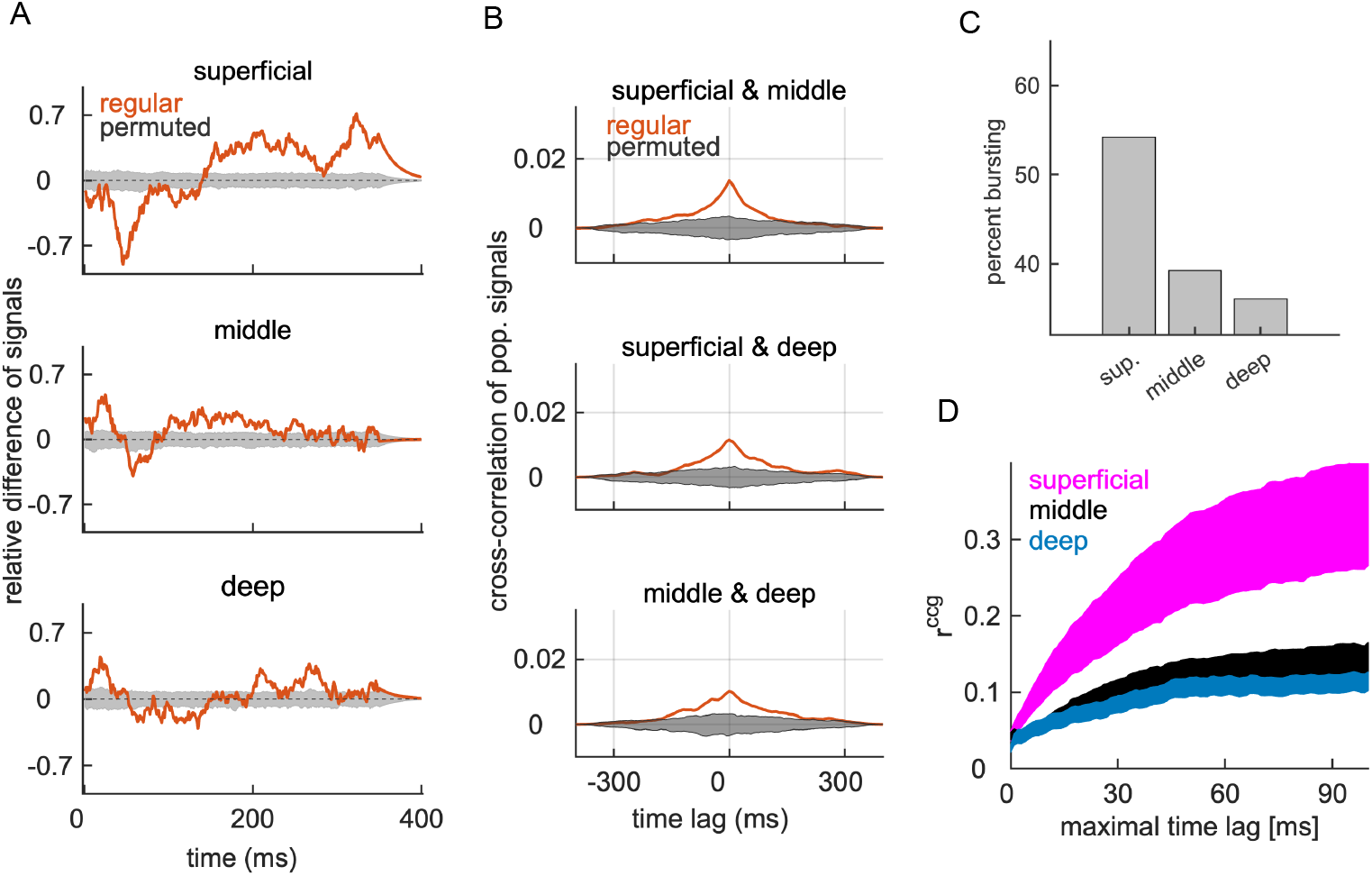
Spiking activity in the superficial cortical layer carries the strongest decision signal. A: Session-averaged difference of decision signals (red) and the distribution of corresponding signals by read-out models with permuted class labels (gray). B: Cross-correlation of decision signals across pairs of cortical layers (red) and the corresponding result of models with permuted class labels (gray area). C: Percentage of bursting versus non-bursting neurons in cortical layers. D: Noise correlation of spike timing for neuronal pairs within specific layers. We show the mean ± SEM across pairs. We used the time window [140,500] ms after the onset of the test stimulus. Parameters: *λ*^-1^ = 20 ms, *N*_perm_ = 1000, time window for A-C is [0, 400], time window for D is [140,500] ms after the onset of the test stimulus.

The superficial layer has an excess of bursting neurons (Fig 7D), which explains stronger signal in the superficial layer compared to other layers. In the superficial layer, 54 % of neurons are bursting neurons (while the remainder are non-bursting). In the middle and deep layer, 39 and 36 % of neurons are bursting, respectively. We have shown above that bursting neurons have stronger correlations than non-bursting neurons. Consistently with the excess of bursting neuron in the superficial layer of the cortex, noise correlations of spike timing are the strongest among neurons in the superficial layer compared to middle and deep cortical layers (Fig 7D).

## Discussion

In the present study, we examined the stimulus- and the choice-related population signals in V1 and predicted the future behavioral choice from the spiking activity in the primary visual cortex. We showed that the choice can be decoded with a time-dependent linear population read-out, and that decoding weights need to be learned in correct trials. We argued that the generalization of learning from the variable stimulus + choice to choice relies on the (partial) overlap in the information on the stimuli and the choice. Furthermore, we found that bursting neurons and neurons in the superficial layer of the cortex are particularly informative for the task. Our results suggested that correlated spiking of neurons with similar decoding selectivity helps the accumulation of the time-resolved choice signal and thus enhances the population signals about the categorical stimulus class and the congruent behavioral choice of the animal.

Current results raise a question, why is learning in correct trials necessary for the prediction of the choice? The following scenario during the condition “incorrect non-match” is compatible with all our observations: 1) the test stimulus is “non-match” and V1 responds with a spiking pattern corresponding to the stimulus class “non-match” during the first 100 or so milliseconds after the onset of the stimulus, 2) there is an error in transmission of the signal to downstream areas that will result in incorrect choice [45], 3) due to the error in transmission, a high-level area erroneously signals the stimulus class “match” and the choice “same” and projects this signal back to V1, and 4) as the projection is received by V1, the spiking pattern in V1 switches and signals the stimulus class “match” and the choice “same”. This scenario allows the activity in V1 to be predictive of the choice simply through the alignment of the representation about the stimuli and the choice, activated by a top-down projection.

If the population signal in the context of stimulus + choice would rely exclusively on the difference in stimuli, we would not be able to predict the variable choice by learning population decoding weights on stimulus + choice. On the contrary, if the representations of the choice and the stimulus class in V1 are mixed [23] and the information about the two variables is overlapping [56], learning in the context of *stimulus* + *choice* can generalize to the representation of *choice*, leveraging the part of information that is invariant across the two classification problems. In the brain, such alignment can be instrumental in the sense that it reinforces the representation of the upcoming behavioral choice across brain networks. The information about the behavioral choice was shown to be distributed across multiple brain areas [57], and alignment of the signals about the stimulus and the choice could serve the purpose of reverberating and amplifying the choice-related signal within the cortical circuitry, to help the brain converge to a single behavioral outcome. Future work could address the overlap of the stimulus and choice-related information with information-theoretic measures [56] that can disentangle the contribution of these sources of information more precisely than the read-out based methods that we used in the present analysis.

Another result that merits the discussion is the role of noise correlations for the time-dependent population signal. Positive noise correlations of spike counts between neurons with similar selectivity are known to decrease the quantity of information in a rate code [51], and they do so also in the present dataset [39]. However, in the present decision-making task, rather than transmitting all the information about the complex naturalistic stimuli, neural networks are faced with the task of extracting task-relevant information (“match/non-match” of stimuli) and creating a variable that can inform the binary choice of the animal. The binary variable related to animal’s choice “same/different” only consists of 1 bit of information. Rather than maximizing the quantity of information about the stimulus, it might be important to create a variable that is robust to trial-to-trial variability of spiking patterns and can be read-out consistently over the course of the trial. Recent theoretical work pointed out that noise correlations among neurons with similar tuning are useful for improving the robustness of a rate code [58] and the temporal consistency of the read-out [53]. In the present work, we show how structured noise correlations can be useful in a time-dependent population read-out. Correlated spiking of neurons with similar decoding selectivity allows accumulation of the population signal and makes the signal rise from the noisy background. Note that if the spiking activity was entirely uncorrelated across neurons, a time-dependent read-out of such uncorrelated spiking would be a noisy oscillation around zero. From such a signal it would indeed be impossible to read-out the choice of the animal or any other behaviorally relevant variable. Present results therefore suggest a simple and biologically relevant role of correlated spiking between neurons with similar selectivity - to allow the accumulation of a time-dependent low-dimensional signal and help its transmission downstream.

How can a top-down signal simultaneously drive neural responses and determine the structure of correlations? Previous work has suggested that the structure of noise correlations in V1 is primarily determined by a common input [16]. Moreover, correlated variability in V1 was shown to change with task instructions [59], strongly suggesting that correlated variability in V1 can be driven by a feedback from higher cortical area(s). In line with these results, we showed that noise correlations are stronger between neurons that contribute the most to the choice signal. As long as top-down projections are specific about the sign of decoding weights of V1 neurons, such projections could at the same time drive the choice signal and create the observed pattern of noise correlations where neurons with similar decoding selectivity are more strongly correlated. Top-down input could target neurons with positive weights in trials with decision “same” and neurons with negative weights in trials with decision “different” through precise long-range connections. Precise long range microcircuit-to-microcircuit connections between the prefrontal cortex and V1, recently put in evidence [60], are a good candidate for conveying such a precise feedback signal. Future work could tell if such a simple mechanism can explain present results.

Finally, our results showed that, in the present behavioral task, bursting neurons are particularly informative about the choice of the animal. Bursting neurons reported here likely correspond to recently reported class of bursting excitatory neurons in the V1 of the macaque, located predominantly in the superficial layers of the cortex [61]. Bursting neurons have been proposed to integrate bottom-up and top-down inputs [62, 63], and be involved in learning of synaptic weights across cortical hierarchy [54]. Encoding of binary stimulus classes and the binary choice is not a build-in feature of V1 neurons and in order for V1 neurons to encode such variables, their synapses with the top-down projecting neurons have to be modified. Bursting neurons might therefore be particularly informative about the choice of the animal in the current experimental setting because their synapses with top-down projecting neurons have been modified in such a way as to support coding of a binary stimulus and choice.

## Conclusion

In the present setting, macaque monkeys learned to classify binary categories “match” vs. “non-match” on succession of two complex naturalistic stimuli. In such a setting, a binary choice variable “same”/“different” in the absence of the information about the stimulus class can be predicted from the spiking activity of neural populations in the primary visual cortex using learning from invariants, a learning method that is likely used in intelligent systems. The time-dependent signal about animal’s decision can be computed as a population read-out of spike trains by the observed neural population and models a variable proportional to the synaptic current in hypothetical read-out neurons. Such a signal carries the information about animal’s choice in decoding weights of bursting neurons in superficial cortical layers, as well as in temporal and across-neuron organization of spike trains. Temporal and across-neuron organization of spike trains is captured by noise correlations of spike timing that seem to be such as to help the accumulation of the decision signal.

## Supporting information

suppose

## Supporting information

**S1 Fig. Robustness of the decision signal with respect to the time window used for decoding.**

**S2 Fig. Learning and validation on the variable** *choice* **result in poor prediction of the decision variable.**

**S3 Fig. Animal’s decision cannot be predicted during the target stimulus.**

## Acknowledgments

This project was supported by Deutsche Forschungsgemeinschaft, grant GRK 1589/2, Technische Universität Berlin and Berlin Equal Opportunity Programm (Berliner Chancengleichheitsprogramm).

