## Supplementary material for "Learning from invariants predicts upcoming behavioral choice from spiking activity in monkey V1": suppose

### Supporting information for "Learning from invariants predicts upcoming behavioral choice from spiking activity in V1"

**S1 Figure: Robustness of the decision signal with respect to the time window used for decoding.**

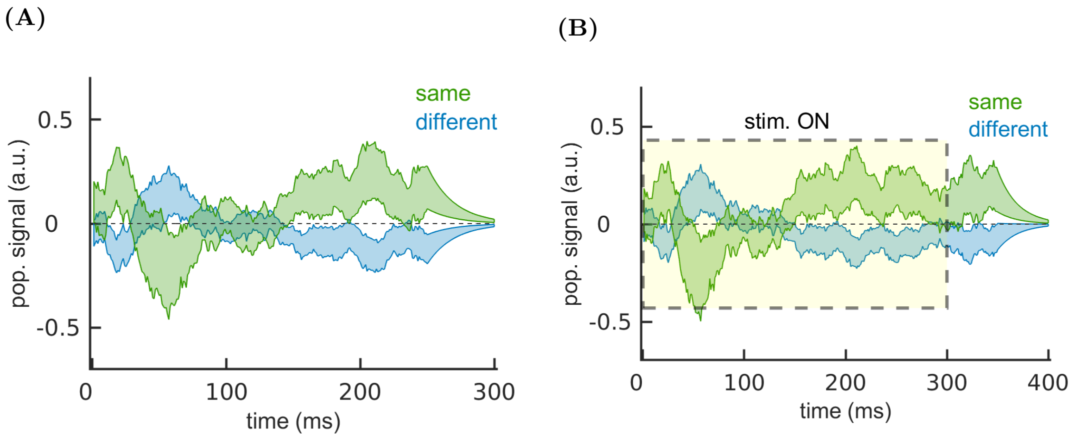


**Figure S1. Robustness of the decision signal with respect to the time window used for decoding.** A: The decision signal computed with the time window $[0, 300]$ ms with respect to the offset of the test stimulus. The same time window is used to estimate decoding weights and to compute the population signal. B: Same as in (A), utilizing the time window of $[0, 400]$ ms respect to the offset of the test stimulus. Parameters: $\lambda^{-1} = 20$ ms, $N_{perm} = 1000$.

**S2 Figure: Learning and validation on the variable *choice* result in poor prediction of the decision variable.**

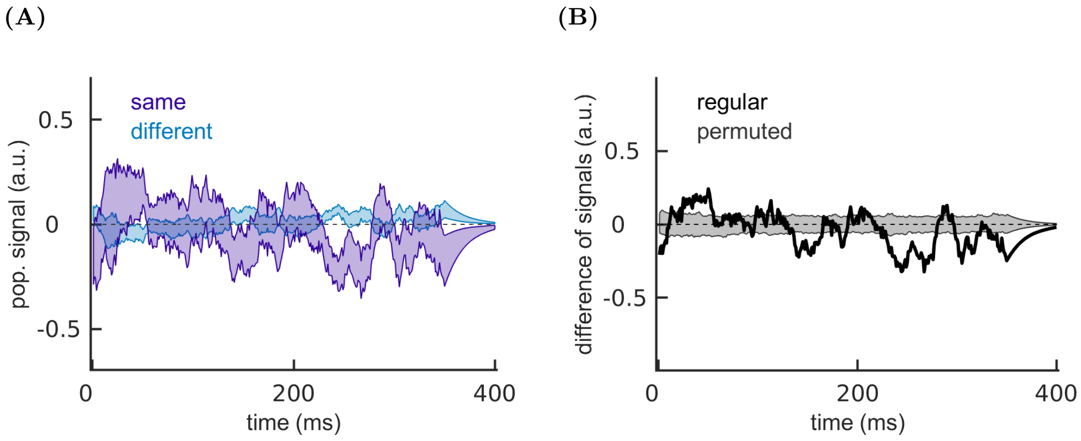


**Figure S2. Learning and validation on the variable *choice* result in poor prediction of the decision variable.** A: Decision signal computed with decoding weights on the variable *choice*. We show the mean ± SEM for the variability across recording sessions. Utilizing the variable *choice* for decoding weights gives noisy signals and poor prediction of the decision variable. B: The difference of mean decision signals across recording session (black) and the distribution of corresponding results for models with permuted class labels. Parameters: $\lambda^{-1} = 20$ ms, $N_{perm} = 1000$.

**S3 Figure: Animal's decision cannot be predicted during the target stimulus.**

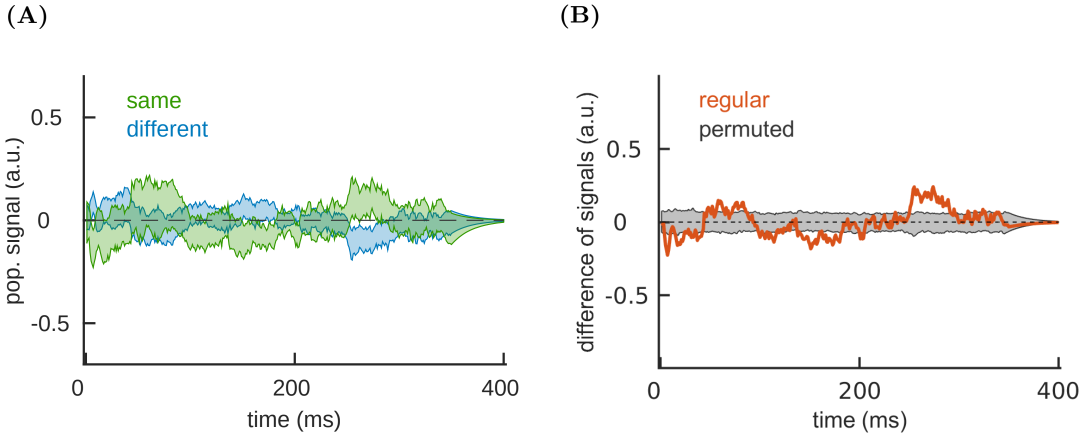


**Figure S3. Animal's decision cannot be predicted during the target stimulus.** A: Decision signal in the time window $[0, 400]$ ms with respect to the onset of the target stimulus. We show the mean $\pm$ SEM for the variability across recording sessions. Neural activity during the presentation of the target stimulus does not predict the choice of the animal. B: The difference of mean decision signals across recording session during the presentation of the target stimulus (black) and the distribution of corresponding results for models with permuted class labels. Parameters: $\lambda^{-1} = 20$ ms, $N_{perm} = 1000$.
